# Strain-specific genome evolution in *Trypanosoma cruzi*, the agent of Chagas disease

**DOI:** 10.1101/2020.07.15.204479

**Authors:** Wei Wang, Duo Peng, Rodrigo P. Baptista, Yiran Li, Jessica C. Kissinger, Rick L. Tarleton

## Abstract

The protozoan *Trypanosoma cruzi* almost invariably establishes life-long infections in humans and other mammals, despite the development of potent host immune responses that constrain parasite numbers. The consistent, decades-long persistence of *T. cruzi* in human hosts arises at least in part from the remarkable level of genetic diversity in multiple families of genes encoding the primary target antigens of anti-parasite immune responses. However, the highly repetitive nature of the genome – largely a result of these same extensive families of genes – have prevented a full understanding of the extent of gene diversity and its maintenance in *T. cruzi*. In this study, we have combined long-read sequencing and proximity ligation mapping to generate very high-quality assemblies of two *T. cruzi* strains representing the apparent ancestral lineages of the species. These assemblies reveal not only the full repertoire of gene family members in the two strains, demonstrating extreme diversity within and between isolates, but also provide evidence of the processes that generate and maintain that diversity, including extensive gene amplification, dispersion of copies throughout the genome and diversification via recombination and *in situ* mutations. These processes also impact genes not required for or involved in immune evasion, creating unique challenges with respect to preserving core genome function while maximizing genetic diversity.

## Introduction

The protozoan parasite *Trypanosoma cruzi* is the causative agent of Chagas disease, the highest impact parasitic infection in the Americas, affecting 10 to 20 million humans and innumerable animals in many species. The study of *T. cruzi* and Chagas disease is particularly challenging for a number of reasons, including the complexity and unique characteristics of its genome. Over 50% of the *T. cruzi* genome is composed of repetitive sequences, which include numerous families of surface proteins (e.g. *trans*-sialidases, mucins and mucin-associated surface proteins) with hundreds to thousands of members each, as well as substantial numbers of transposable elements, microsatellites and simple tandem repeats (Weston et al. 1999; El-Sayed et al. 2005; De Pablos and Osuna 2012). This repetitive nature greatly hampered the assembly of the original CL Brener strain reference genome generated in 2005, resulting in a highly fragmented and draft assembly with extensively collapsed high repeat regions (El-Sayed et al. 2005). In addition, the CL Brener strain turned out to be a hybrid strain with divergent alleles at many loci. To scaffold the genome sequence, Weatherly *et al*. took advantage of the bacterial artificial chromosome (BAC) library sequencing data and combined with synteny analysis of two genomes from closely related species, *Trypanosoma brucei* and *Leishmania*, obtained the current reference genome with 41 chromosomes (Weatherly et al. 2009). Nevertheless, a large number of gaps are still present in the chromosomes of the reference genome, and many unassigned contigs remain, making it impossible to determine the exact genome content and, in particular, the full repertoires of large gene families.

As in many pathogens, and best documented in the related trypanosomatid *Trypanosoma brucei*, families of variant surface proteins often serve as both the primary molecular interface with mammalian hosts and as the predominant target of host immune responses. Classical antigenic variation in these pathogens consists of the serial expression of a single (or highly restricted number of) antigen variant(s) in the pathogen population at any one time, with switches to new variants becoming evident once the host immune response controls the dominant one. This largely “one-at-a-time” strategy appears particularly effective in pathogens exposed continuously to antibody-mediated immune control mechanisms. *T. cruzi*, however, appears to take a much different approach to antigenic variation, generating multiple very large families of genes encoding surface and secreted proteins, many of which are expressed simultaneously rather than serially. We believe that this strategy may reflect the primarily intracellular lifestyle of *T. cruzi* in mammalian hosts and the necessity of evading T cell recognition of infected host cells, although this has yet not been formally proven.

The advent of two advances in genome analysis has made it feasible to revisit and substantially improve upon the *T. cruzi* genome assembly and to advance our understanding of its composition. The long-read capability of PacBio Single-Molecule Real-Time (SMRT) sequencing provides read lengths capable of spanning long repetitive regions. The application of this technology (Berna et al. 2018; Callejas-Hernandez et al. 2018) as well as nanopore sequencing (Diaz-Viraque et al. 2019) has resulted in much-improved contiguity and expansion of gene family members in *T. cruzi*. Secondly, proximity ligation methods have allowed for the scaffolding of assemblies spanning highly repetitive regions. One of the methods, Hi-C, identifies extant inter-chromosomal interactions by capturing chromosome conformation, and has been used to create scaffolds at chromosomal scale (Kaplan and Dekker 2013; Korbel and Lee 2013). A second approach termed Chicago, adapts this same methodology but reconstitutes the confirmation of DNA *in vitro* by combining the DNA with purified histones and chromatin assembly factors (Putnam et al. 2016). These proximity ligation methods not only improve the contiguity of genomes by joining contigs, they also identify misjoins in the contigs and separate them to increase the accuracy of assemblies (Putnam et al. 2016). The combination of Chicago and Hi-C has now been applied to many genomes (Robert D. Denton 2018; Theodore S. Kalbfleisch 2018; Elbers et al. 2019; Salter et al. 2019; Schreiber et al. 2020).

In this study, we have applied SMRT sequencing and proximity ligation methods to produce very high-quality assemblies from the Brazil (TcI) and Y (TcII) strains of *T. cruzi*. These two strains are representatives of the most ancestral lines that are hypothesized to have given rise to the 6 discrete typing units (DTUs, TcI-TcVI) lineages now composing this genetically diverse species (Westenberger et al. 2005; de Freitas et al. 2006; Zingales et al. 2009; Flores-Lopez and Machado 2011; Zingales et al. 2012; Tomasini and Diosque 2015). Using these chromosomal-level assemblies with minimal gaps, we are now able to compare the full gene content of representatives of these founding lineages of the *T. cruzi* species, including the full repertoires of large gene families. Herein, we document a substantial diversity in individual chromosome content, including frequent allelic variants, but with an overall conserved gene content outside of the large gene families. Within these gene families, however, extreme diversification is evident with no genes of the identical sequence within strains or shared between these strains. These high-quality genomes also reveal the mechanisms behind the expansion and diversification of the large gene families, presumably in response to immunological pressure, and in the process, creating other challenges in terms of core genome stability and function.

## Results

### Genome Sequencing and Assembly

PacBio SMRT sequencing provided 1,264,527 (N50=9,560 bp) and 763,579 (N50=12,499 bp) filtered reads with ∼9 Gb and ∼6 Gb of sequence data for Brazil clone A4 (Brail A4) and Y clone C6 (Y C6), respectively, corresponding to ∼200x and ∼130x coverage based on the predicted genome size. Initial assembly resulted in sequences of 45.11 Mb and 46.98 Mb for Brazil A4 and Y C6 draft genomes, respectively, close to the estimated haploid genome size of *T. cruzi* (Souza et al. 2011) (Table 1). Application of the *in vitro* proximity-ligation tools Hi-C and Chicago (Putnam et al. 2016), decreased the L50 to half of that of the draft genomes, and the size of the largest scaffolds doubled (Table 1). Filling gaps and base correction using Illumina reads ultimately resulted in 12 and 14 scaffolds in the Brazil A4 and Y C6 final assemblies, respectively, with a length greater than 1 Mb. Telomeric repeats [(TTAGGG)n] were identified in 18 Brazil A4 and 15 Y C6 scaffolds, including on both ends of three scaffolds in Brazil A4, suggesting full chromosome assembly in these cases. The improvement in these new genomes is not only in integrity (Supplemental Table S1 and Supplemental Fig. S1), but also in filled gaps, recovered genes and extended repetitive regions (see examples in Supplemental Fig. S2).

**Table 1.**
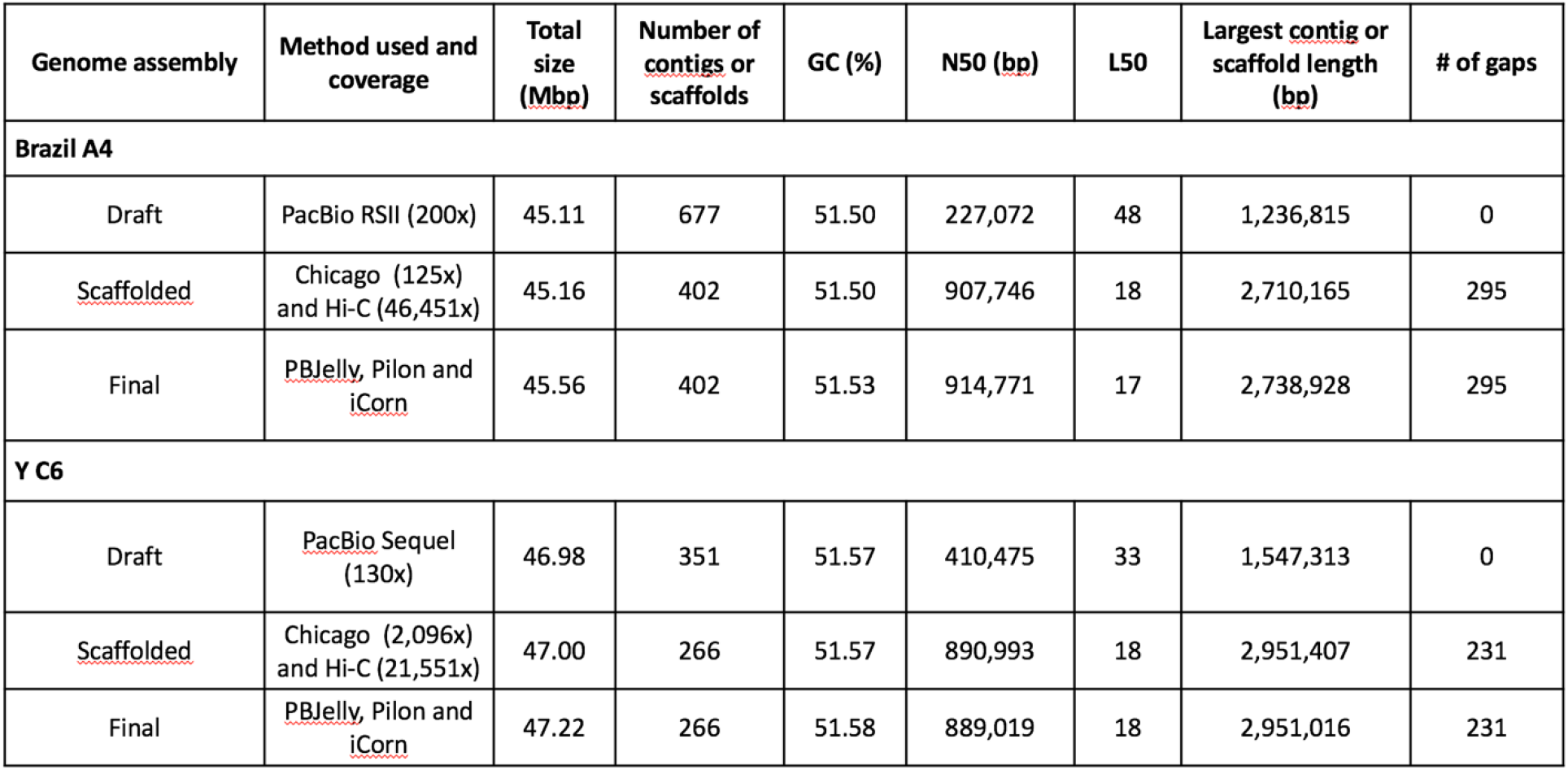
Summary of assembly statistics.

### Genome features and content

Due to a lack of apparent chromosome condensation during replication (Henriksson et al. 2002; Souza et al. 2011), the karyotype of *T. cruzi* has not been completely elucidated. Moreover, chromosome size and content vary significantly between different *T. cruzi* strains and even among clones of the same strain based upon pulse-filed gel electrophoresis (PFGE) analysis (Henriksson et al. 2002; Pedroso et al. 2003; Vargas et al. 2004; Triana et al. 2006; Lima et al. 2013). Based on criteria including size, repeat proportion, and gene number, 43 scaffolds of Brazil A4 and 40 scaffolds of Y C6 were designated as chromosomes (Supplemental Fig. S3) and the remainder referred to as smaller scaffolds.

Repetitive sequences occupy 58.8% and 62.3% of the genome for Brazil A4 and Y C6 (Supplemental Tables S2 and S3), substantially higher than the 50% that was estimated in the reference CL Brener genome, thus confirming the capability of long-read sequencing and assembly approaches to recover and place more repetitive DNA content. Approximately 50% of the sequence in chromosomes is repetitive sequences, compared to ∼90% in smaller scaffolds (Supplemental Fig. S3). Using conventional approaches with manual curation, gene models were identified in Brazil A4 and Y C6, respectively. Based on BUSCO assessment, the Brazil A4 and Y C6 contain the highest number of single-copy gene sets among assembled *T. cruzi* genomes (Supplemental Table S4).

A major constituent of the repetitive regions in the *T. cruzi* genome is large gene families, including the *trans*-sialidases (TS), mucin associated surface proteins (MASP), mucins, and surface protease GP63 (all targets of immune responses), as well as retrotransposon hotspot (RHS) proteins and dispersed gene family 1 proteins (DGF-1) (El-Sayed et al. 2005; Buscaglia et al. 2006; Martin et al. 2006). Our previous studies indicated the total copy number of TS genes was underestimated using conventional annotation approaches due in part to the failure to identify new variants and fragments of TS resulting from frequent recombination (Weatherly et al. 2016). To complete the annotation of the members of large gene families, we developed a customized workflow (summarized in Supplemental Fig. S4) and applied it to the six largest gene families. This allowed us to capture the full repertoire of gene family members (copy numbers of which are summarized in Supplemental Table S5), the distribution of which were plotted in Fig. 1 and Supplemental Fig. S5. Gene family members are unequally distributed among and along the chromosomes with several of the largest chromosomes (e.g. TcBrA4_Chr2 and TcYC6_Chr1) composed nearly entirely of gene family members. In contrast to previous reports suggesting the members of large gene families were mainly located in telomeric and subtelomeric regions (Carlos Talavera-López 2018) (El-Sayed et al. 2005), gene family members are not restricted to particular regions of chromosomes. Moreover, TS, MASP, mucin and GP63 have an overlapping distribution along the chromosomes, while RHS and DGF-1 genes are more dispersed.

**Figure 1.**
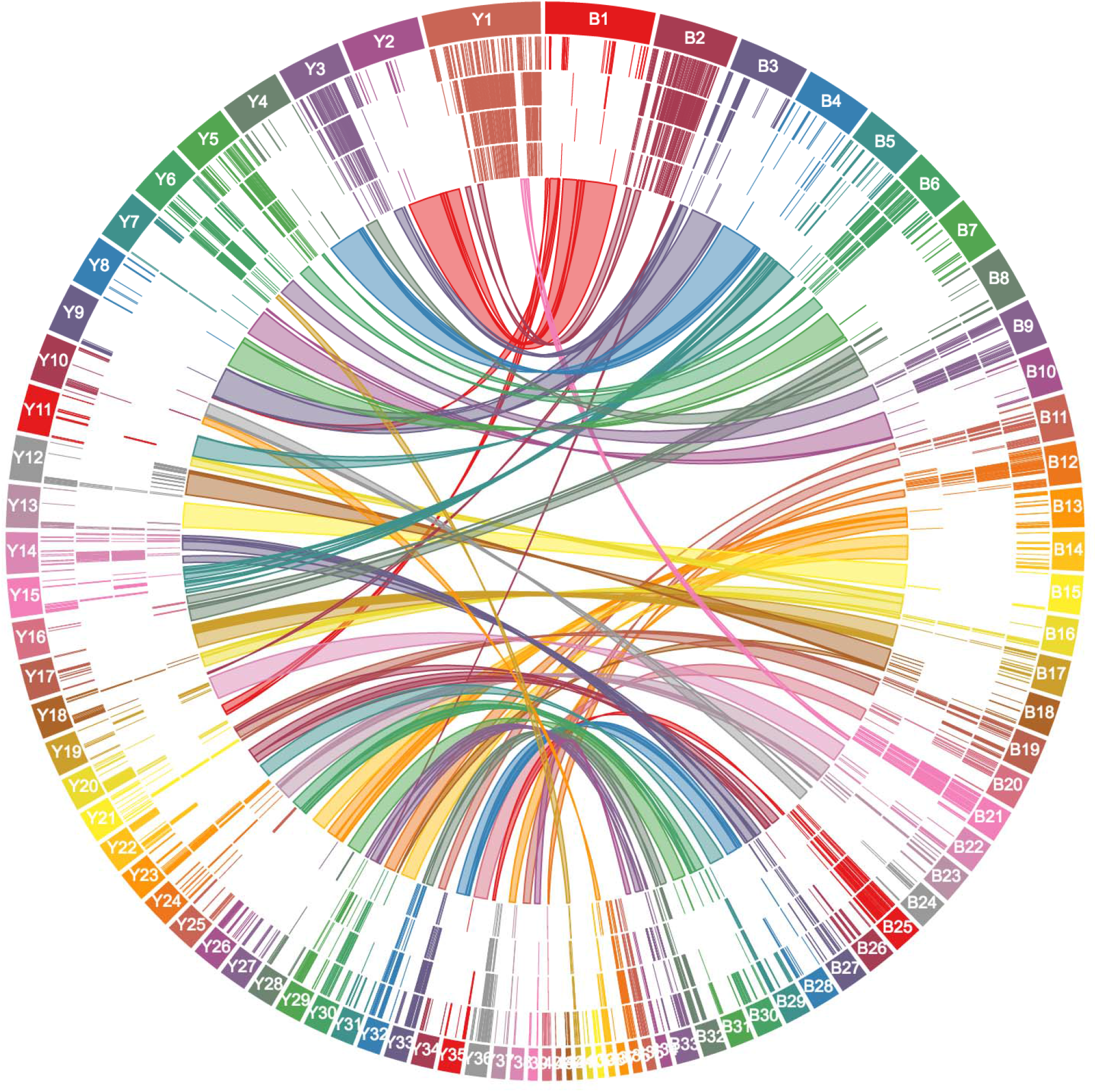
Distribution of large gene families and synteny between chromosomes in Brazil A4 (right, B) and Y C6, (left, Y). Tracks from outer to inner rings: chromosomes, TS, MASP, mucin, GP63, and synteny blocks.

After consolidating the predictions of large gene families with our conventional annotations, the Brazil A4 and Y C6 genomes contained 18,708 and 17,650 gene models, respectively (see annotation summary in Supplemental Table S6). The composition of gene content between two genomes is very similar, with ∼25% as members of large gene families, ∼40% as hypothetical proteins, and >90% of the remaining genes are orthologs of those in the related kinetoplastids *T. brucei* and *Leishmania major*. That this gene model count in the two *T. cruzi* strains is substantially higher than that estimated for *T. brucei* and *L. major* is likely due to two factors: 1) the high number of large gene family members in *T. cruzi*, and 2) a greater number of hypothetical genes in *T. cruzi*, a third of which are unique to *T. cruzi*, although the size distribution of the hypothetical proteins is similar in the 3 species (Supplemental Fig. S6).

### Allelic variation

The significant number of small scaffolds and the relatively high gene model numbers in some of the small scaffolds prompted us to consider whether these small scaffolds might represent regions of allelic variation between sister chromosomes, as allelic variation is one of the factors that results in fragmentation during genome assembly for diploid genomes. Although TcI and TcII DTUs represented by the Brazil and Y strains, respectively, are considered homozygous lineages, we very conservatively detected 26 and 33 small scaffolds in each genome showing consistent synteny in multiple gene models to parts of the core chromosomes (Supplemental Table S7). An example is shown in Fig. 2A in which scaffold TcYC6_Contig191 demonstrates regions of synteny within the 825,025 – 1,379,353 bp region in the first chromosome of Y C6 (TcYC6_Chr1). Confirmation of this chromosome variant was supplied by replacing the identified region in TcYC6_Chr1 with TcYC6_Contig191 and then mapping the chromosomal contacts in the Hi-C data for these 2 alternative versions for TcYC6-Chr1. As shown in Fig. 2B, the Hi-C data are equally strong for both chromosome variants. Using Falcon-Phase, which phases diploid genome sequences by integrating long reads and Hi-C data (Zev N. Kronenberg 2018), we identified an additional 18 and 7 allelic variations in Brazil A4 and Y C6, respectively. In combination, these analyses identified allelic variations in 24 chromosomes of Brazil A4 and 25 of Y C6, including chromosomes with multiple allelic variants, e.g. the largest chromosome in Y C6 (TcYC6_Chr1), and an intermediate-sized chromosome in Brazil A4 (TcBrA4_Chr13; Fig. 2C). Thus, we suggest that many of the small scaffolds are variants of regions in the chromosome-size scaffolds. However, because the majority of these small scaffolds lack the conserved, non-gene family sequences required to prove synteny, and Falcon-Phase can only resolve haplotypes bearing divergence of < 5%, identifying the position of all the small scaffolds on the chromosomes was not possible.

**Figure 2.**
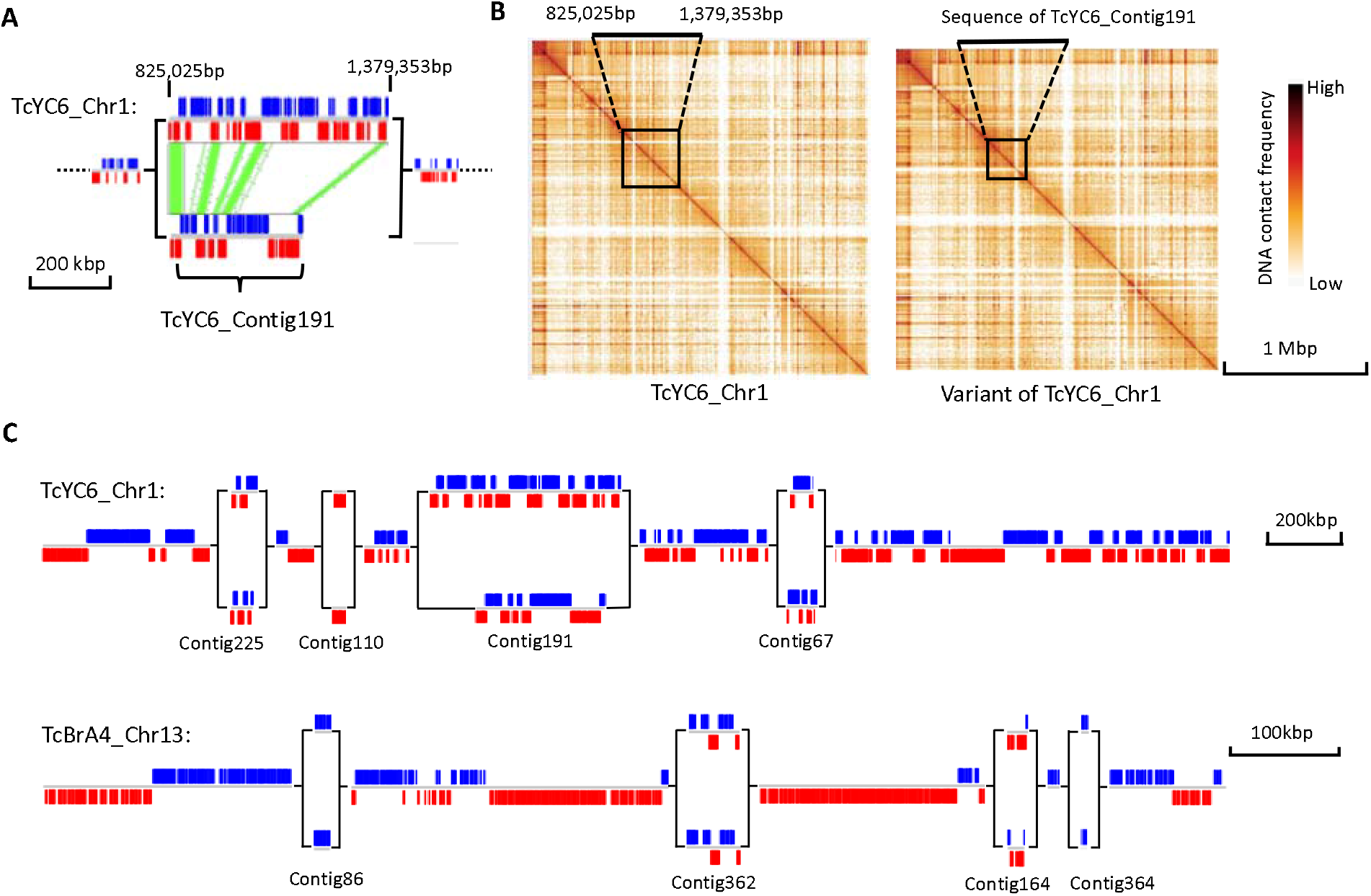
An example of homologous chromosomes with large allelic variations. (A) Synteny between two allelic variants in Chr1 of Y C6. Synteny blocked are marked with green. (B) Hi-C heat maps of TcYC6_Chr1 (left) and its homologous chromosome with TcYC6_Contig191 (boxed area) replacing the allelic region in TcYC6_Chr1 (boxed area). (C) Two chromosomes with multiple allelic variants. Blue blocks indicate genes on the forward strand, and red blocks indicate genes at the reverse strand.

### Structural comparison of the Brazil and Y sequences

The very high genome quality and contiguity provided by the combination of SMRT sequencing and Hi-C analysis enabled chromosome level comparison of the Brazil (TcI) and Y (TcII) clones (Fig. 1). The synteny plots show that the majority of chromosomes from one genome collinear with those in the other genome. For instance, Brazil A4 Chr4 showed continuous synteny to Y C6 Chr4 overall. However, as expected based upon previous gene mapping studies (Henriksson J 1990) (Henriksson et al. 1995) (Vargas et al. 2004) (CaroleBranchea 2006), some chromosomes corresponded to different regions in multiple chromosomes of the other genome, e.g. Brazil A4 Chr1 showed synteny to a combination of Chr20 (298,235 - 684,393bp), Chr9 (63,384 - 95,053bp) and Chr2 (20,327 - 1,438,658bp) in Y C6. Some inverted syntenies were also detected, e.g. between 388,900 - 968,190bp on Brazil A4 Chr8 and 11,711 - 556,982bp on Y C6 Chr16 (Fig. 1). Notably, the diversity of sequences encoding members of the large gene families (see details below) prevented the detection of synteny in a substantial proportion of the two genomes, including in two of the largest chromosomes (e.g. TcBrA4_Chr2 and TcYC6_Chr1).

### Variation in gene models within and between Brazil and Y strains is predominantly in the large gene families

A large number of genetic variations were identified in the non-repetitive region, including heterozygous SNPs/Indels within respective strains, and homozygous SNPs/Indels between the two strains (Supplemental Tables S8, S9 and S10). We also detected aneuploidy in both genomes: 3 and 8 chromosomes in Brazil A4 and Y C6, respectively, exist in copy numbers greater than two, based on the results of both relative read depth and allele frequency (Supplemental Fig. S7). Among these are the partially syntenic chromosomes (TcBrA4_Chr24 and TcYC6_Chr10), which also share synteny with chromosome 31 in CL Brener, reported to be supernumerary in many strains (Reis-Cunha et al. 2015), thus suggesting a species-wide requirement for > 2 copies of one or more genes in these regions. Additionally, variation exist in the copy number for a substantial number of individual genes characterized by OrthoFinder, with ∼150 genes showing the greatest variation between the two strains (Supplemental Table S11). However, with respect to genes unique to either strain, we found 23 (Brazil A4) and 20 (Y C6) gene loci not present in the other strain and further validated this finding by examining the raw reads (Supplemental Table S12). All are annotated as hypothetical proteins and most are small genes located in gene family-rich regions of the genome and thus are likely the products of recombination events involved in gene family diversification (see below).

To fully assess the variation in the large gene family members between the two strains, we carried out a best match search for the protein sequence of putatively expressed genes in each large gene family from Y C6 genome with those in Brazil A4. As a control, the same analysis was performed for a subset of mostly single-copy genes (BUSCO), as well as a small gene family of 35 members, beta galactofuranosyl glycosyltransferase (b-gal GT). As shown in Fig. 3, high-identity matches could always be found for the BUSCO genes, and some of them (22 out of 291) have identical matches (100% identity) in the other strain. Similar to BUSCO genes, the identity between best matches for b-gal GT is also tightly distributed in the range of 90-97%. In contrast, all large gene families exhibit a broad distribution of identity for their best matches relative to the BUSCO genes and b-gal GT genes, especially TS, MASP, mucin and RHS, with only a small proportion of best matches bearing 90% identity or more. Among the family members with the greatest similarity between the two strains are the small subset of TS genes containing the sialidase enzymatic domain as previously described (Cremona et al. 1995), suggesting that this group of *trans*-sialidases has been selected for and conserved in both strains.

**Figure 3.**
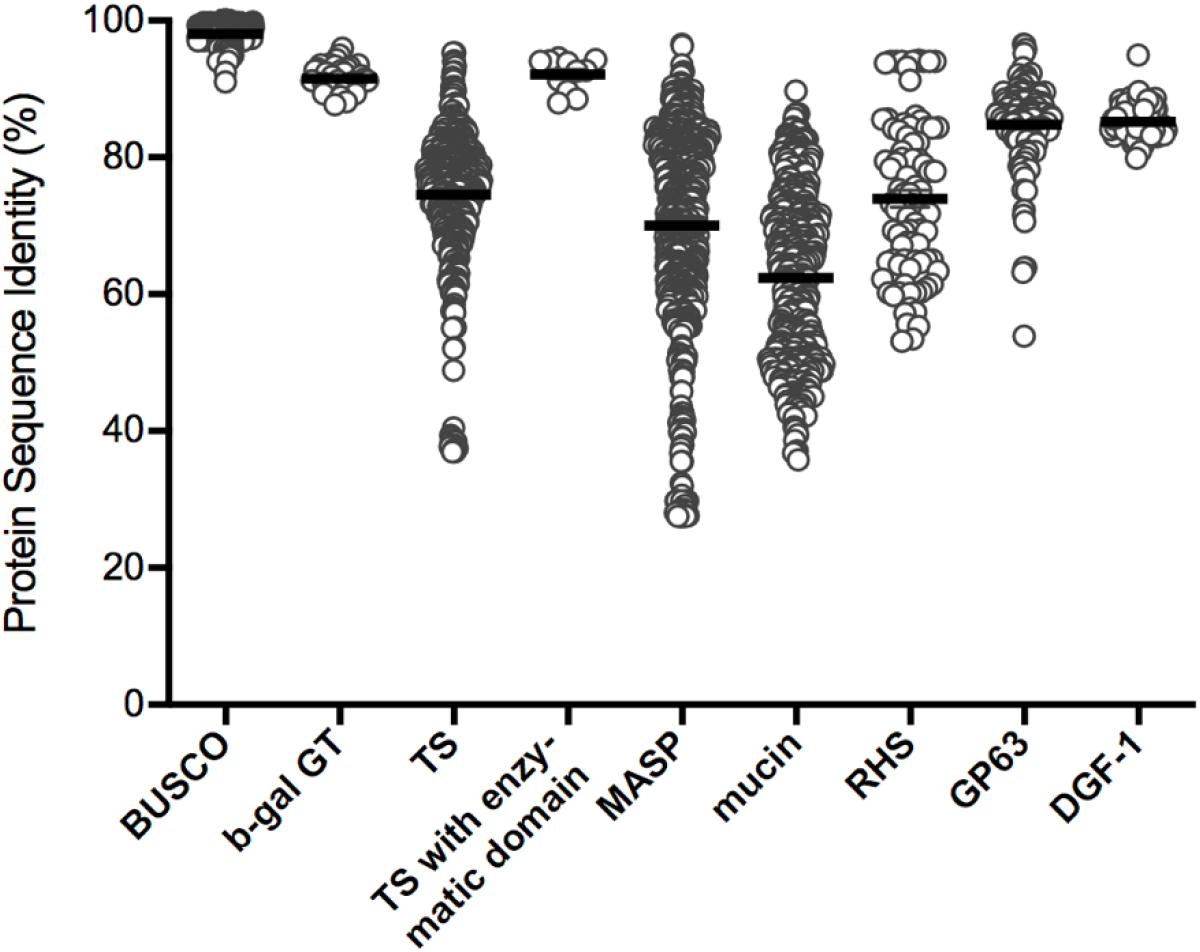
Protein best match analysis of gene families between Brazil A4 and Y C6.

### Evidence of gene family expansion and diversification

The very high number and the impressive within- and between-strain variation in the genes composing the large gene families in *T. cruzi* is indicative of a system under intense evolutionary pressure. We have taken advantage of the high contiguity of these two genome sequences, as well as the comprehensive prediction of all members of large gene families, to attempt to understand better how this remarkable diversity is generated and maintained.

We first examined the genomes for evidence of gene duplication events that could increase the number of members in gene families. Multidimensional scaling (MDS) plots based on the pair-wise genetic distances of all members of each gene family in each strain allowed us to identify tightly distributed gene clusters with high sequence identity (http://shiny.ctegd.uga.edu). In multiple cases, genes within these clusters were tandemly arrayed individually (TS; Fig. 4A top) or as a set of genes (TS plus MASP; Fig. 4A bottom). Such tandem amplifications are present in all gene families (except DGF-1) and occur uniquely in each strain (Supplemental Table S14). A number of unusual amplification events were also noted, including inverted duplications creating a strand switch in between (Fig. 4B), and an amplification involving several genes on both strands, replicated a total of 5 times (Fig. 4C), thus creating a complex set of strand switches.

**Figure 4.**
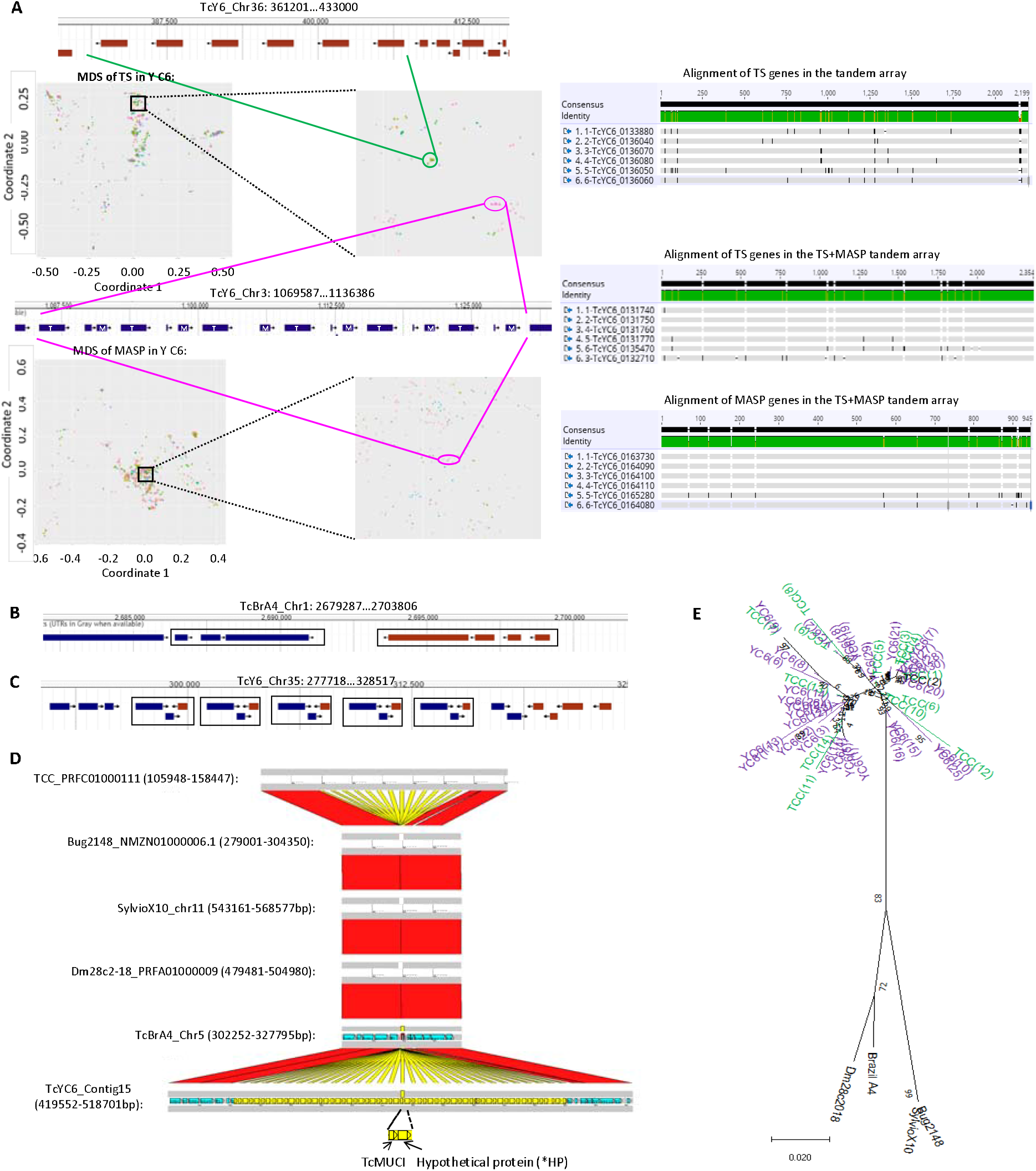
Gene amplification events in members of large gene families. (A) Tandem arrays of individual TS genes (top), and a TS+MASP pair (bottom) clustered based upon genetic distance in the MDS plots. Each chromosome is displayed as a separate pattern on the MDS plot. T: TS; M: MASP. Alignment of the genes in each MDS cluster (right) confirms high consensus (grey regions); black regions indicate SNPs and ‘–’ indicate gaps. (B) Mirror-duplication of one fragmented RHS and two pseudo RHS genes. (C) One RHS (+), one hypothetical protein (+) and one fragmented glycosyltransferase (-) replicated 5 times, creating multiple strand switches. (D) Syntenic regions of the TcMUCI+*HP tandem array detected in 6 long-read sequenced *T. cruzi* strains. Synteny of TcMUCI orthologs are labeled in yellow. (E) Phylogenetic tree of all TcMUCI orthologs from the 6 strains. Note that TcMUCI genes from Y C6 (purple) and TCC (green) are intermingled in the top portion of the tree. Live MDS plots can be explored at http://shiny.ctegd.uga.edu.

The majority of tandem amplification of gene families in both *T. cruzi* genomes contained 10 or fewer replicates (Supplemental Table S14). However, one hypothetical protein (*HP) in the Y C6 occurr in a tandem array of 29 units with a TcMUCI gene (Fig. 4D). Comparison to the syntenic region in Brazil A4 revealed a single TcMUCI ortholog (and no *HP sequence), indicating that at some point the *HP sequence was inserted next to the TcMUCI gene in Y C6, and the two genes were amplified together as a segment (Fig. 4D). Although no particular protein domains were characterized in the *HP gene, 18% of its sequence share similarity with several MASP sequences, implying that at least part of the gene might be derived from a MASP. The abnormally high number of replicates in this tandem array as well as the low diversity in the tandem copies suggest that this might be a recent amplification event. However, comparison to syntenic regions in other long-read sequenced *T. cruzi* genomes revealed the same TcMUCI+*HP tandem array in the TCC (TcIV) strain but not in the Dm28c (TcI), Sylvio (TcI), and Bug2148 (TcV) (Fig. 4D). Additionally, phylogenetic analysis grouped all of the replicated copies from Y C6 and TCC together and distant from the single TcMUCI genes in the other three strains (Fig. 4E). Using the model of DTU evolution in *T. cruzi* which postulates that the TcVI is derived from a hybridization event between TcII and TcIII (Westenberger et al. 2005; Tomasini and Diosque 2015), we propose that the TcMUCI+*HP amplification is an ancient event, occurring after the split of TcI and TcII but prior to the TcII/TcIII hybridization that yielded TcVI.

In addition to tandem clusters of gene family members, MDS analysis also revealed closely related gene family members located on multiple chromosomes (Fig. 5A). An extreme case is the Y C6 gene cluster in the bottom right of Fig. 5A which contained 31 TS genes with very high similarity distributed on 11 different chromosomes (Supplemental Fig. S8A). Interestingly, the 13 TS on Chr7 (Fig. 5B, middle) are in tandem, interspersed with a beta tubulin gene, while the 7 TS on Chr18 (Fig. 5B, top) are in tandem as TS genes alone (with one beta tubulin gene downstream of TS_7_). The remaining 11 TS genes in this cluster are dispersed in the genome as singlets (3 of them are shown at the bottom of Fig. 5B). Notably, the sequences upstream and downstream of the TS gene in the TS + beta tubulin array on Chr7 are homologous to those of the TS_7_ gene in the Chr18 array, and the dispersed singlet TS also share a portion of the upstream and downstream sequences with the other TS in this cluster (Supplemental Fig. S8B). Together, these results suggest that all 31 TS genes in this cluster originated from one or more gene amplification/relocation events. Based on the phylogenetic analysis (Supplemental Fig. S8C), we propose that the TS + beta tubulin tandem copies have been generated in or relocated to Chr7 (13 copies) and Chr18 (1 copy), with another 4 TS copies as single genes beyond the TS + beta tubulin cassette on Chr18, while the single TS genes on other chromosomes may derive from TS on Chr7.

**Figure 5.**
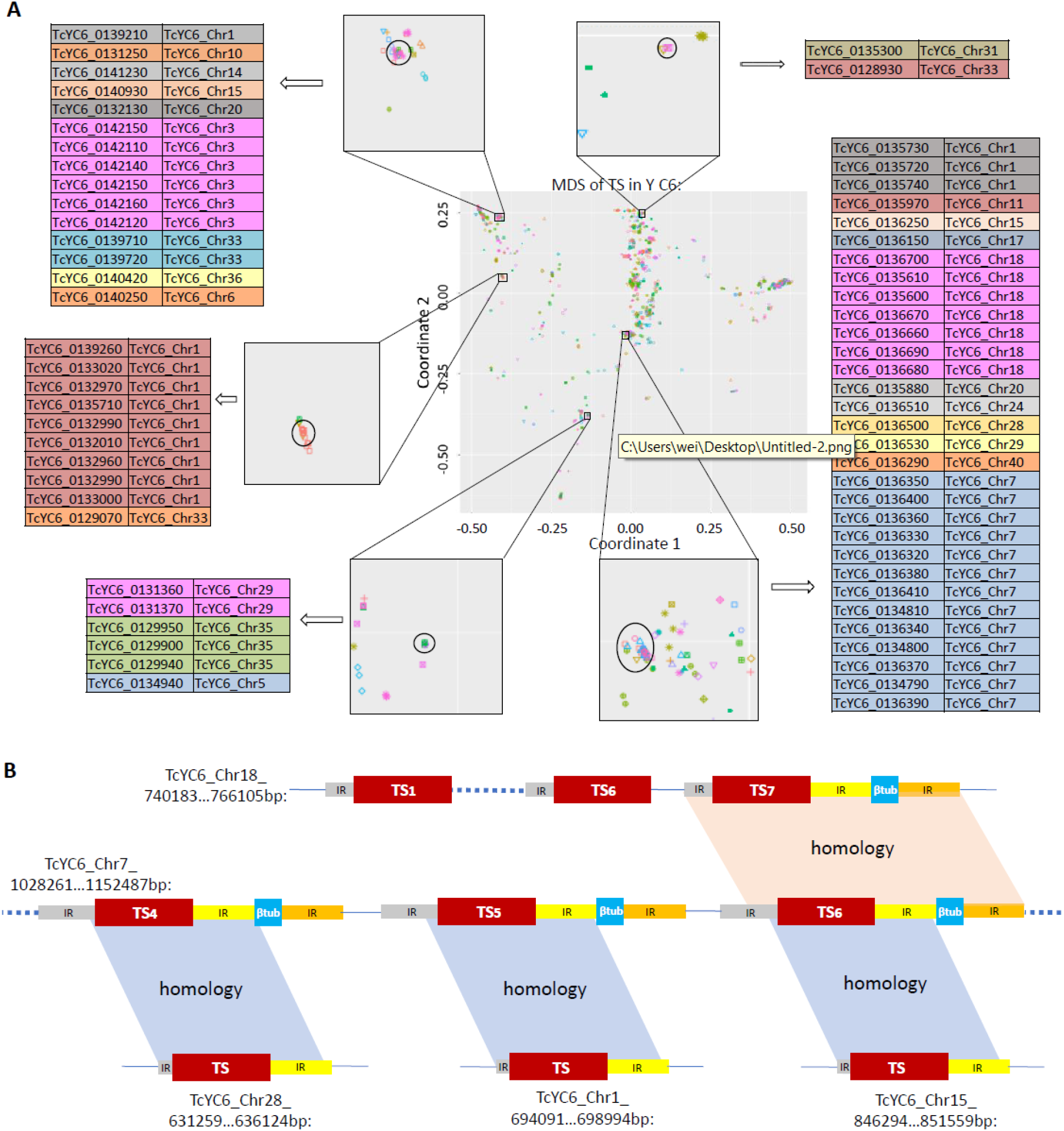
Examples of relocations of TS genes in Y C6. (A) Tight clusters of TS genes from MDS plot are distributed on different chromosomes. (B) Diagram of relocations in one of the TS clusters on the bottom right in (A). Blocks in the same color indicate genes or flanking sequences in high identity. IR: intergenic region. Note that the segment size is not to scale.

We next used a pipeline previously designed to identify recombination events within TS genes in the CL Brener genome (Weatherly et al. 2016), to quantify recombination for 4 of the large gene families in the Brazil and Y strains (Table 2). As expected, recombination events, including multiple events acting on the same gene, were detected in a large fraction of the genes but were particularly abundant (2-fold higher) in the TS family relative to the other three families examined. Interestingly, recombination events in the TS family were detected at a roughly 2-fold higher frequency in the Brazil strain as compared to the Y or the CL Brener strains.

**Table 2.** Recombination events detected within genes of large gene families in Brazil A4 and Y C6.

|  | Brazil A4 |  |  |  | Y C6 |  |  |  | CL Brener |
| --- | --- | --- | --- | --- | --- | --- | --- | --- | --- |
|  | TS | MASP | Mucin | GP63 | TS | MASP | Mucin | GP63 | TS |
| # of genes | 1644 | 1118 | 700 | 411 | 1465 | 1066 | 797 | 427 | 3209 |
| Kb length total | 3477.7 | 1011.9 | 352.1 | 460.6 | 2614.5 | 1115.2 | 458.9 | 619.7 | 4456.5 |
| # of genes recombined | 793 | 145 | 38 | 70 | 479 | 154 | 73 | 89 | 787 |
| # of recombination events | 2976 | 190 | 39 | 101 | 1334 | 221 | 85 | 153 | 2087 |
| % of genes recombined | 48.2 | 13.0 | 5.4 | 17.0 | 32.7 | 14.4 | 9.2 | 20.8 | 24.5 |
| Average events per gene | 1.8 | 0.2 | 0.1 | 0.2 | 0.9 | 0.2 | 0.1 | 0.4 | 0.7 |
| Average events per kb | 0.9 | 0.2 | 0.1 | 0.2 | 0.5 | 0.2 | 0.2 | 0.2 | 0.5 |
| Number of genes with n recombination events |  |  |  |  |  |  |  |  |  |
| n=1 | 137 | 111 | 37 | 51 | 162 | 110 | 61 | 58 | 324 |
| n=2 | 198 | 24 | 1 | 15 | 114 | 28 | 12 | 19 | 149 |
| n=3 | 98 | 9 | 0 | 1 | 66 | 11 | 0 | 3 | 110 |
| n=4 | 128 | 1 | 0 | 1 | 54 | 3 | 0 | 6 | 72 |
| n=5 | 68 | 0 | 0 | 1 | 33 | 2 | 0 | 1 | 52 |
| n>5 | 164 | 0 | 0 | 1 | 50 | 0 | 0 | 2 | 80 |
| Max of n | 18 | 4 | 2 | 5 | 12 | 5 | 2 | 6 | 12 |

As noted previously, our recombination pipeline is highly conservative in detecting relatively recent events that have not been obscured by subsequent accumulation of SNPs and Indels (Weatherly et al. 2016). Such *in situ* diversification is evident in genes that are clustered in the MDS analysis but dispersed in the genome. An example is a cluster of GP63 genes in Brazil A4 which have low genetic distance based on MDS analysis (Supplemental Fig. S9A), but are located on different chromosomes and display a considerable degree of variation (SNPs and Indels; Supplemental Fig. S9B). However, because these genes also share similar upstream genes (a TS) and intergenic regions, all of these dispersed genes were likely derived via gene duplication. This hypothesis is further supported by the result that 9 out of 10 GP63 genes and their corresponding GP63 + flanking sequences (including upstream TS + intergenic region + GP63 + intergenic region) occupy identical positions in their respective phylogenetic trees (Supplemental Fig. S9C). Therefore, a TS/GP63 gene pair and associated intergenic regions underwent one or more duplication and relocation events with subsequent diversification through the accumulation of SNPs and Indels, yielding multiple, diverse genes spread through the genome.

The potential complexity generated by amplification, relocation, recombination and diversification make it challenging to track the specific set of events contributing to the evolution of individual gene family members in *T. cruzi*. However, some gene sets reveal all of these processes at work. Fig. 6C shows four cassettes located on different chromosomes or in distant sites on the same chromosome, each cassette with a central MASP and flanking region with high identity (Fig. 6A and B), suggesting a common origin. SNPs/Indels indicate *in situ* diversification of the MASP genes, especially in the C terminus (Fig. 6B). Cassette pairs I/II and III/IV share the same upstream gene and flanking sequence (mucin genes in both cases) while cassette pairs I /III and II/IV shared downstream mucin and GP63 genes, respectively. In addition, a recombination event was detected in the C terminus of the GP63 in cassette IV, creating divergence from the GP63 C terminus in cassette IV.

**Figure 6.**
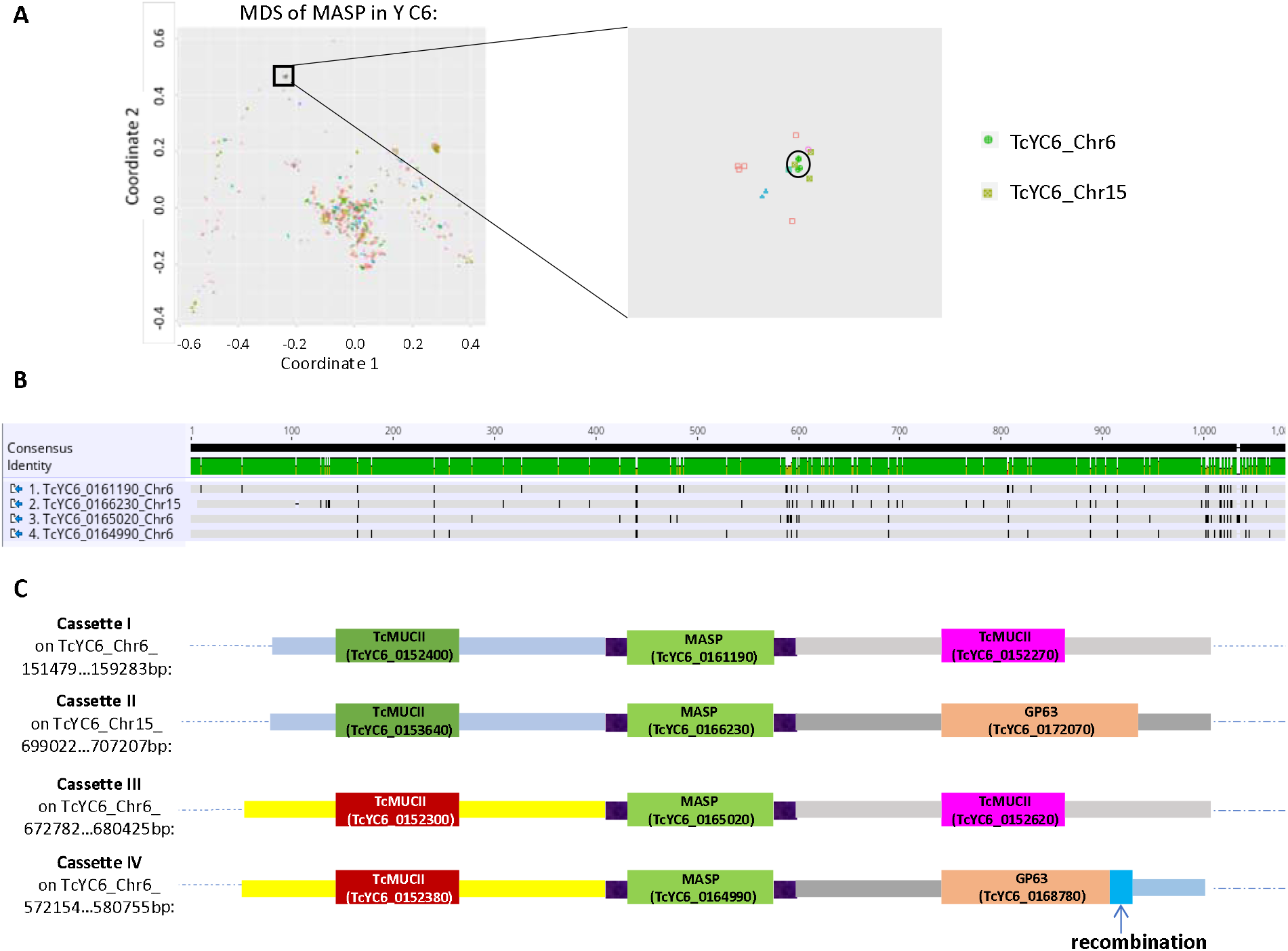
The combination of gene amlification, relocation, recombination and *in situ* diversification of large gene family members. (A) A tight cluster of 4 MASP genes from MDS plot are distributed on two chromosomes. (B) Alignment of the 4 MASP genes shows high identity with modest diversification of SNPs/Indels. (C) MASP genes with flanking intergenic sequences and flanking genes. Blocks with the same color indicate sequences in high identity. Note that the segment sizes are not to scale.

### Potential impact of high genome flexibility on gene expression

Unlike in other classical models of antigenic variation in protozoa, the large gene families in *T. cruzi* are not restricted to particular regions of chromosomes [e.g. subtelomeric in the case of *T. brucei* (El-Sayed et al. 2003; Berriman et al. 2005; Hertz-Fowler et al. 2008; Mugnier et al. 2016; Muller et al. 2018)] but instead are spread throughout the genome (Fig. 1). This presents the complication that the amplification and dispersion events common in the large gene families of *T. cruzi* might also impact non-gene family (core) genes as well. To investigate this possibility, we focused on core genes for which there were > 6 total paralogues for the two genomes, and organized these paralogues on the basis of gene location (Supplemental Table S11). By doing this, we could identify tandemly distributed genes that likely resulted from gene amplification. For the over 150 groups of genes in this analysis, many showed dramatic differences in gene copies in the two *T. cruzi* genomes with 26 instances of double-digit gene copies in one strain compared to only 1-3 copies in the other. This same high level of variation was also evident for other *T. cruzi* genomes sequenced using long-read sequencing methods but not in similarly sequenced *T. brucei* and *Leishmania* isolates (Supplemental Fig. S10). Additionally, dispersion patterns for these amplified genes differed widely between the Y C6 and Brazil A4 genomes (Supplemental Table S11). Thus, the mechanisms that provide for the generation and maintenance of diversity in the large gene families also appear to allow for substantial variation in copy number for selected core genes, and representing a second major contributor to between-strain genetic variation in *T. cruzi* strains.

Most gene expression in trypanosomatids initiates in the absence of specific promoters and with the production of multi-gene mRNA transcripts that are then processed into single-gene mature mRNAs. These polycistronic transcriptional units (PTUs) of genes can be well over >100 kb in length and are marked by start and stop signals, including base modifications (Clayton 2019). The apparent wide degree of freedom for amplification and dispersion both within and outside the *T. cruzi* large gene families, and particularly events that create tandem strand switches as shown in Fig 4C, would be expected to impact this normal multi-gene PTU structure. Indeed, the average PTU length was ∼116.5 and 126.8 kb in the core gene-rich regions of the Brazil A4 and Y C6, respectively, similar to that in *T. brucei* (148.3 kb). However, the average PTU length in the gene-family-enriched regions of both *T. cruzi* genomes was less than ¼ of that (29.3 kb in Brazil A4 and 33.8 kb in Y C6), indicating a disruption of the normal PTU structure. Interestingly, amplified but conserved tandem gene arrays like the ‘mucin + *HP’ array in Y C6 discussed above (Fig. 4D) and the previously described TcSMUG family (Yoshida 2006; Nakayasu et al. 2009; Gonzalez et al. 2013) are within large PTUs containing almost no gene family members (Supplemental Fig. S11A) while many other tandem arrays or apparently diverging genes reside in gene-family-rich, short PTUs (Supplemental Fig. S11B). The disruption in PTU structure might also hamper the preservation of transcriptional control mechanisms, in particular the tight controls on transcriptional termination characterized in other kinetoplastids and mediated by base J and histone H3/4 variants (Siegel et al. 2009; Cliffe et al. 2010; van Luenen et al. 2012; Reynolds et al. 2014; Reynolds et al. 2016; Schulz et al. 2016; Kawasaki et al. 2017; Muller et al. 2018). To address this question, we mapped strand-specific RNA-seq reads to both sense and antisense strands to assess transcriptional termination relative to PTUs. Surprisingly, we found extensive antisense RNA levels throughout the genome (an average of sense:antisense=114:1 in Brazil A4 and 84:1 in Y C6). Higher levels of antisense RNAs occurred at the strand switch regions of long PTUs (Fig.7A and B), but in some cases, matched or exceeded the sense strand transcripts in gene-family-enriched regions containing shorter PTUs (Fig. 7C). Thus, unlike *T. brucei* and *Leishmania, T. cruzi* does not appear to regulate antisense RNA production so tightly.

**Figure 7.**
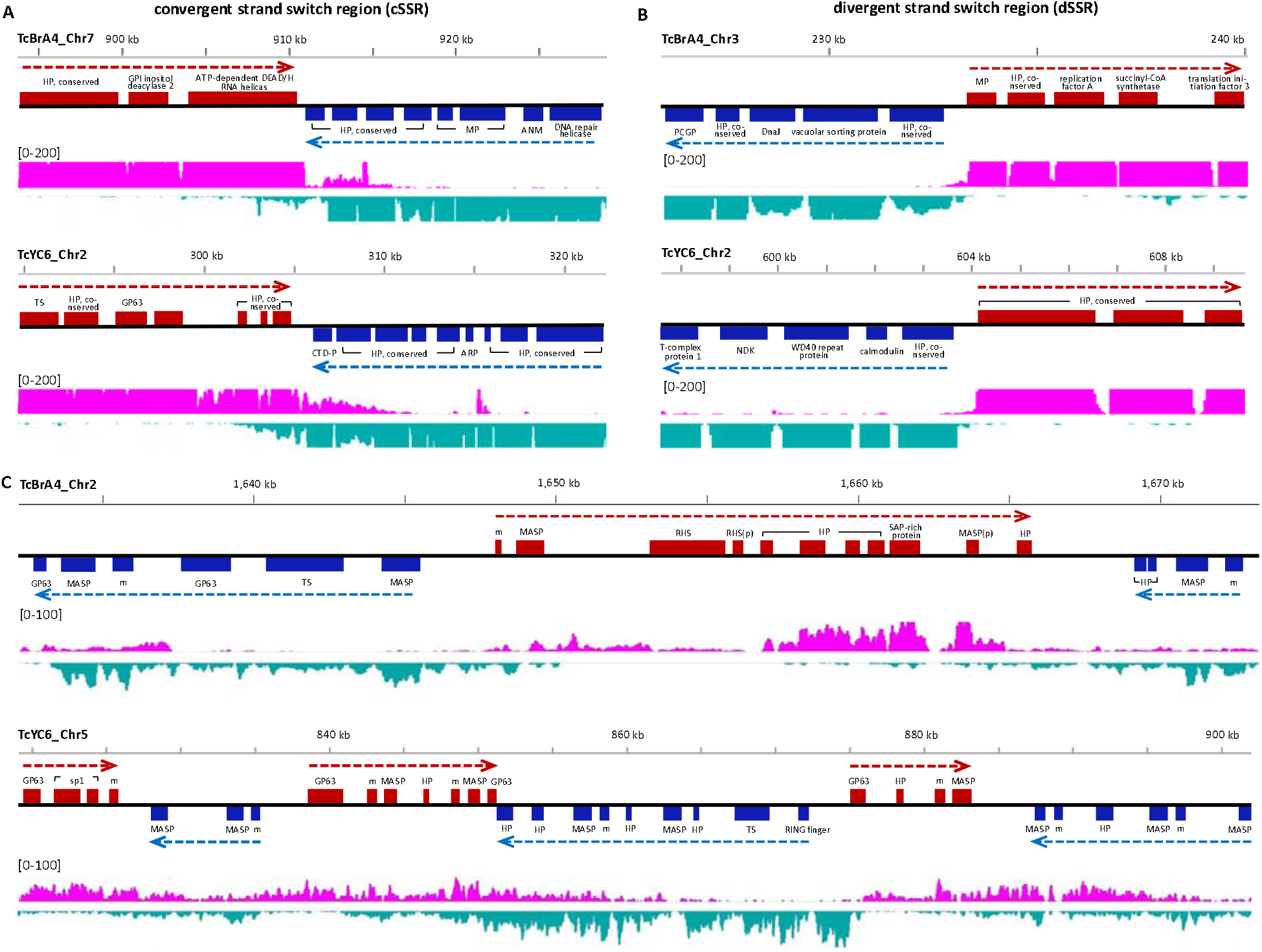
Antisense RNA levels in *T. cruzi* in relation to PTU structure, including both convergent strand switch regions (cSSR, A) and divergent strand switch regions (dSSR, B). (C) Gene-family-enriched regions with frequent strand switches where antisense RNA were detected in higher levels. HP, hypothetical protein; MP, mitochondrial protein; ANM, arginine N-methyltransferase; PCGP, parkin coregulated gene protein; CTD-P, TFIIF-stimulated CTD phosphatase; ARP, ankyrin repeat protein; NDK, nucleoside diphosphate kinase; SAP-rich protein, serine-alanine- and proline-rich protein; m, mucin; p, pseudogene.

## Discussion

*T. cruzi* is a highly heterogeneous specie, with at least six DTUs and with extreme variation in phenotype and virulence among isolates even of the same DTU. Gaining new understanding of the genetic basis of this high strain-to-strain variation in disease-causing potential in *T. cruzi* has been challenging due to the lack of high-quality reference genome. The high content of repetitive sequences in the *T. cruzi* genome (>50%), including multiple families of surface protein-encoding genes each with >200 members, makes complete genome assembly from conventional short-read sequences impossible. This study reports very high-quality genomes for *T. cruzi* strains belonging to the presumed ancestral lineages of this species, TcI, represented here by the Brazil strain and TcII by the Y strain. This significantly improved resource was achieved by the application of long-read sequencing techniques and proximity ligation libraries to better resolve the full repertoires of gene content, thus allowing a detailed comparison of genetic variation between these strains.

Although DTU-specific associations have been frequently proposed for characteristics such as virulence, disease presentation, geographic distribution, and host species restrictions, many of these linkages falter when more extensive sampling is done and none has been linked to DTU-specific genetic differences (Revollo et al. 1998; Rassi et al. 2012; Nguyen and Waseem 2020). The current dataset provides the opportunity to begin examination of representative strains of *T. cruzi* lineages that diverged from each other an estimated 1-3 million years ago (Tomasini and Diosque 2015). The most surprising revelations from this comparative analysis were not the variability in unique gene content between these isolates, but rather the extremes of the high similarity in core gene content and the comparative huge diversity in gene family-rich portions of the genomes. As anticipated based upon previous strain-based screens (Ackermann et al. 2009) (Reis-Cunha et al. 2015), a considerable degree of variation exists in the form of SNPs/Indels and additionally, a substantial number of strain-specific copy number differences were identified. However, the core (non-gene family) genome, contains only ∼20 strain-unique gene models, and in all cases, these are hypothetical genes encoding proteins with no recognizable protein domain structures.

In very sharp contrast, the variation evident in the large gene families of *T. cruzi* is equally remarkable, demonstrating vast diversity within and between strains with no perfect matches and relatively few genes of the same family with even a 90% similarity. Structurally, these gene family members make up ∼25% of the genome and are spread widely throughout the genome, with some members on every chromosome and some of the largest chromosomes being almost entirely composed of gene family members. The use of synteny detection tools and Falcon-Phase validated by Hi-C methods allowed us to also conservatively document heterozygosity in more than half of the chromosomes in each genome, and we suspect that this heterozygosity extends to nearly all gene family-rich regions of the genome. Based upon the total base count of the repeat-rich small scaffolds not assigned to chromosomes, we estimate that up to 50% of all gene family members have variants on the sister chromosome.

The quality of the genome assemblies also provided the opportunity to document the continuing diversification of these large gene families and to permit the beginning of an understanding of how this process might work. Select members of the large gene families in *T. cruzi* have clear and critical functions in parasite biology, with the best-documented example being the enzyme-active *trans*-sialidases required for acquisition of sialic acid by *T. cruzi* trypomastigotes (Previato 1985) (Uemura 1992; Cremona et al. 1995) (Frasch 2000). However, the number, diversity, and potential for variation of genes in these large gene families, and the exposure of the gene products to and response by the host immune system, argue that these gene families evolve under intense immunological pressure. In this respect, the three largest and most diverse gene families in *T. cruzi* (TS, MASP and mucin) are similar to other families of genes involved in antigenic variation in the protozoans *T. brucei* (variant surface glycoproteins, VSGs), *Plasmodium* (*var* genes) and Giardia (Variant-specific Surface Protein, VSPs) (Cross 1975; Mowatt et al. 1991; Pimenta et al. 1991; Smith et al. 1995; Su et al. 1995). However in contrast to the “one-at-a-time” models of classical antigenic variation best characterized in the sister kinetoplastid *Trypanosoma brucei* (Cross 1975), *T. cruzi* expresses many gene family variants simultaneously. This difference in strategy may relate to the fact that *T. cruzi* lives predominantly intracellularly in mammals and must effectively evade cell-mediated (rather than exclusively antibody-mediated) immunity. But expressing many antigen variants at one time also likely requires a larger antigen repertoir and/or an enhanced ability to generate new variants. In African trypanosomes, a comparison of the genome sequences of 2 different **subspecies** (*T. brucei brucei* vs *T. brucei gambienese*) revealed that >86% of the genes – including most VSGs – varied by <1% between the subspecies and only 69 ortholog pairs (including 35 VSG gene pairs) had less than 95% nucleotide identify (Jackson et al. 2010). The diversity of antigen variants in *T. cruzi* among the 2 **strains** examined in this study is vastly greater, with the average similarity of orthologues pairs ranging from 62.4% to 84.8% in the 5 largest gene families (Fig. 2). This finding suggests high pressure to generate variants and a genetic system that accommodate the genomic flexibility that such generation would require.

Classically, segmental duplication creates the source material on which mutational and recombinational events act to derive new genes and new gene functions (Lynch and Conery 2000). The presence of segmental duplications (one gene or multiple genes as a unit) also encourages additional rounds of duplications that can rapidly change gene content (Sturtevant 1925; Muller 1936; Lewis 1951). These processes of gene duplication, recombination and mutation-driven diversification, functioning in concert to ensure high and constant antigenic diversity, is strongly evident in the large gene families of *T. cruzi*. Although we are able to track a significant number of these events, all occurring independently in these two *T. cruzi* strains, we are presumably only observing the most recent occurrences, as recombinations and mutations ultimately obscure the origins of new genes. Certainly the repeat-rich structure and dense representation of retrotranposons of the *T. cruzi* genome facilitates maintenance of these processes and the dispersion of gene family members throughout the genome, and interestingly not restricted to chromosomes ends as is the case in *T. brucei* (El-Sayed et al. 2003) (Berriman et al. 2005) (Hertz-Fowler et al. 2008) (Mugnier et al. 2016) (Muller et al. 2018). However, the specific structural elements that initially established and continue to allow for these apparently constant rearrangements throughout the genome but without impacting overall genome integrity, remain unidentified. From our analysis, no consistent pattern of structures, such as the A/T tracks associated with gene application events in *Plasmodium* (Huckaby et al. 2019) were evident.

The apparent high frequency and continued evolution of gene families in *T. cruzi* also create structures and products unique among the kinetoplastids, including the lack of segregation of gene families to chromosome ends, absence of partitioning of expression sites [as in *T. brucei* VSGs (Ersfeld et al. 1999) Navarro and Gull 2001; (Navarro and Gull 2001; Hertz-Fowler et al. 2008)], tolerance for the generation of short PTUs and frequent strand switching, and most surprisingly, the tolerance of antisense RNA production. The latter may well explain the absence of the machinery for RNAi in *T. cruzi* (DaRocha et al. 2004) (Barnes et al. 2012). The presence of abundant and nearly genome-wide antisense RNAs also suggests that *T. cruzi* does not adhere to the full set of rules for transcription termination as defined in *T. brucei* and *Leishmania* (Reynolds et al. 2014) (Kieft et al. 2020).

Interestingly, there are several subsets of gene family members that appear to be exceptions to these processes of recombination, diversification and distribution throughout the genome. The previously characterized SMUG families are the best examples. Two subgroups of TcSMUG genes, TcSMUG L and S, involved in development and infectivity of insect-dwelling stages (Yoshida 2006; Nakayasu et al. 2009; Gonzalez et al. 2013), distribute as tandem arrays in the respective subgroups within the same PTU and exhibit minimal diversification. Here we also identify an ancient, lineage-specific duplication event that created a new hypothetical gene and a mucin gene in a tandem array and which, like the SMUGS, has remained with minimal changes. It will be of interest to determine if further diversification of this and other gene family subsets are restricted because of their location in the genome, or if, like the SMUGS, this hypothetical gene/mucin tandem is under selective pressure due to their unique function. One common feature of these tandem arrays is that they all locate in and are flanked by large PTUs (>220 kb) containing only core genes with no members from the large gene families (other than the mucins in the mucin+*HP array), suggesting that they are maintained in an environment largely devoid of large-gene-family-related diversification.

An additional strain-dependent difference documented here is the higher recombination frequency in Brazil A4 compared to Y and in CL Brener (Weatherly et al. 2016). The ∼2X greater number of recombination events in all gene families in Brazil vs Y suggests that this is an inherent property of this strain and perhaps of DTUI strains in general. Alternatively, because we very conservatively call recombination events which then eventually become concealed by further mutations/recombinations over time, it is also possible that the Brazil A4 has been under stronger, or more recent, strong selective pressure.

The apparent high levels of gene amplification/diversification readily documented in the large gene families in this species also extends to a fraction of core genes as well, and represents a second major source of between-strain diversity and perhaps the one primarily responsible for the broad between-strain phenotypic variation in *T. cruzi*. Retention of these core gene amplifications imply a fitness benefit, perhaps under certain environmental/host conditions; others may also occur regularly but engender a fitness cost and thus are lost.

In summary, the careful analysis of these two *T. cruzi* strains soundly confirms the vast genetic diversity of parasite lines within this species, and identifies the bulk of diversity to be represented in 3 compartments: 1) rapidly evolving families of genes involved in immune evasion, 2) a subset of “core” genes not linked to evasion but which vary greatly in copy number and perhaps expression, and 3) SNPs and Indels common to all genomes. We hypothesize that the gene family diversity is driven by immune selection and that the same processes that provide for this diversity also allow for copy number variation and diversification of select core genes, and this later process, rather than DTU type, accounts for much of the biological diversity of *T. cruzi* lines. With these high quality genomes in hand for these strains, we can now test these hypotheses by further modifying these gene sets and exposing both wild-type and modified parasite lines to various levels of selection pressure and observing the genomes of the lineages that emerge.

## Supplementary files

**Supplemental Figure S1**. Overview of the Brazil A4 and Y C6 genomes. Tracks from outer to inner circles indicate: sizes, chromosomes, gaps, gene density (window size: 20kb, range: 6-23 in Brazil A4, 1-22 for Y C6), GC content (window size: 10kb, range: 0.36-0.70 in Brazil A4, 0.33-0.70 in Y C6), repetitive content (window size: 10kb, range: 0-10000), heterozygous SNPs (window size: 20kb, range: 0-120 in Brazil A4, 1-390 in Y C6) and heterozygous Indels (window size: 20kb, range: 65-1 in Brazil A4, 129-1 in Y C6).

**Supplemental Figure S2**. Assembly improvement compared to CL Brener. (A) An example of filled gaps. Syntenic regions between Chr1 in Brazil A4 and Chr8 in CL Brener were aligned with the Artemis Comparison Tool (ACT) (Carver et al. 2005). All five gaps were filled in Brazil A4. (B) An example of recovered genes. Two pieces of an adenosine monophosphate (AMP) gene were identified flanking a gap, while the syntenic region in Brazil A4 shows the intact AMP gene. (C) An example of extended repeats. With 8 copies of histone H4 in Chr2 of CL Brener separated by a gap, the syntenic region of Brazil had the gap filled, extending the copy number of histone H4 to 41. Blocks: gaps; green bars: genes.

**Supplemental Figure S3**. Repetitive composition in the scaffolds. Chromosomes are calculated individually, while small scaffolds are calculated by averaging a range of scaffolds as indicated on the x axis.

**Supplemental Figure S4**. Workflow of predicting full repertoire of large gene families (taking TS as an example).

**Supplemental Figure S5**. Distribution of large gene families and retrotransposons on the chromosomes. Rings from outer to inner: chromosomes, retrotransposons, TS, MASP, mucin, GP63, RHS and DGF-1 gene families.

**Supplemental Figure S6**. Size distribution of hypothetical proteins identified in kinetoplastids. Genomes of *T. brucei* TRE92 and *L. major* Friedlin were downloaded from TritrypDB database (https://tritrypdb.org/tritrypdb/) release-44 (Aslett et al. 2010).

**Supplemental Figure S7**. Analysis of chromosome copy number. (A) Relative read depth of each chromosome normalized to the mean read depth of all chromosomes at non-repetitive regions. Chromosomes with more than two copies was indicated in red. (B) Allele frequency calculated by the proportion of heterozygous SNPs/Indels at the non-repetitive regions of each chromosome. A diploid chromosome showed the peak of allele frequency around 50% as shown in chr8 and chr31, whereas a multi-ploid chromosome showed peak of allele frequency lower than 50% as shown in chr24 and chr28 in Brazil A4. Note that 5 chromosomes (Chr35, 36, 38, 39 and 42) in Brazil A4 were not included in this analysis due to their high proportion of repetitive features.

**Supplemental Figure S8**. Alignment of TS (A) and their flanking regions (B) of the cluster in Figure 5, as well as the phylogenetic tree of all the TS genes (C).

**Supplemental Figure S9**. An example of *in situ* diversification. (A) A tight cluster of GP63 genes from MDS plot are distributed in different chromosomes. (B) Alignment of these GP63 genes showed high identity as well as a number of diversifications including SNPs and Indels. (C) Phylogenetic trees of GP63 in the cluster (left), and GP63 plus flanking sequences on both sides (right).

**Supplemental Figure S10**. Copy numbers of 152 orthologue genes sets from Supplemental Table S11 are highly correlated (Spearman correlation > 0.7) in pairwise comparisons between *T. brucei* strains and subspecies and between *Leishmania* species, but poorly correlated between *T. cruzi* strains (range 0.006 - 0.6).

**Supplemental Figure S11**. Examples of long or short PTUs with tandem gene arrays. (A) Tandem arrays of conserved gene sets are contained within long PTUs devoid of gene family members. Chromosomes containing TcSMUG S/L in Brazil A4 (top) and Y C6 (middle), and TcMUCI+*HP in Y C6 (bottom). In contrast, large gene family members, including some tandemly duplicated genes, show frequent strand switches and are in short PTUs (B). Blue bars indicate genes other than large gene family members, while yellow bars indicate large gene families.

**Supplemental Table S1**. Evaluation of assembly metrics among all available *T. cruzi* genomes assembled by long-read sequencing. *No scaffolding was applied to these genomes, so no gaps were generated. **47 are not *de novo* assembled contigs or scaffolds, but rather pseudomolecules produced by aligning the core regions of scaffolds to the core regions of CL Brener reference genome. Therefore, although the, genome showed higher N50 and lower L50, it left an extensively high number of gaps behind. ***Genome sequence is not available.

**Supplemental Table S2**. Repetitive sequences characterized in Brazil A4.

**Supplemental Table S3**. Repetitive sequences characterized in Y C6.

**Supplemental Table S4**. BUSCO assessment of gene completeness for *T. cruzi* genomes with annotation available. Not that TCC is a hybrid strain, so its genome is a mixture of two haplotypes, while all other genomes contain one haplotype.

**Supplemental Table S5**. Copy number of large gene families characterized in the new genomes.

**Supplemental Table S6**. Annotation summary.

**Supplemental Table S7**. Scaffolds that were detected to be allelic variants. Syntenies were examined between small scaffolds and chromosomes. Only those with multiple syntenic regions throughout the entire scaffold with part of the chromosome were considered as allelic variants.

**Supplemental Table S8**. Heterozygous SNPs/Indels identified in Brazil A4.

**Supplemental Table S9**. Heterozygous SNPs/Indels identified in Y C6.

**Supplemental Table S10**. Homozygous SNPs/Indels identified between Brazil A4 and Y C6.

**Supplemental Table S11**. Orthologue groups in *T. cruzi, T. brucei* and *Leishmania* species with total gene count > 6. All sequences were retrieved from TritrypDB database (https://tritrypdb.org/tritrypdb/) release-44.

**Supplemental Table S12**. List of unique genes in the respective strains.

**Supplemental Table S13**. BLAST result of the best match analysis in 6 large gene families between the two strains.

**Supplemental Table S14**. Prominent tandem arrays of large gene families identified in Brazil A4 and Y C6.

## Methods

### Parasite cultures, DNA/RNA extraction and sequencing

Epimastigotes of Brazil and Y were cultured at 26°C in supplemented liver digested-neutralized tryptose (LDNT) medium as described previously (Xu et al. 2009). Single-cell clones were made for each strain by depositing epimastigotes into a 96-well plate at a density of 0.5 cell/well by using a MoFlow cell sorter (Dako-Cytomation, Denmark). One healthy clone that has confirmed to have cycled through all life stages was chosen for sequencing for each strain. High molecular weight DNA was isolated using MagAttract HMW DNA kit (Qiagen) before submitting to Duke Center for Genomic and Computational Biology (GCB) for SMRT sequencing. Brazil A4 was sequenced using PacBio RS II sequencer, while PacBio Sequel sequencer was used for Y C6.

Genomic DNA of the selected clone of both strains was isolated using QIAamp DNA blood mini kit (Qiagen) for whole genome sequencing using Illumina HiSeq 150 PE. An RNase treatment step was included to eliminate RNA in the samples. For RNA-seq sampling, extracellular amastigotes and trypomastigotes isolated from infected Vero cells were pooled with epimastigotes for total RNA-extraction. Following ribo-depleted RNA library construction and RNA sequencing using Illumina Nextseq 75PE was performed by Georgia Genomics and Bioinformatics Core (GGBC). Illumina reads from either DNA or RNA sequencing with mean quality lower than 30 (Phred Score based) were removed for analysis.

### Genome assembly

The draft genome of Brazil A4 was assembled with SMRT Link v3.1, and Y C6 with SMRT Link v5.0. The parameters were set at default except the expected genome size, which was set at 40 Mb for both strains. Chicago and Hi-C libraries were constructed and sequenced by Dovetail Genomics, and HiRise pipeline was run for scaffolding the draft assembly by incorporating data from both libraries. A gap was generated whenever two contigs were joined or one contig was broken by HiRise and since the distance between two contigs was unknown, all gaps were given 100 Ns.

Gap filling was performed by PBJelly (English et al. 2012) using the SMRT subreads with the minimum percent identity at 85%. 122 and 4 gaps were extended for Brazil and Y, respectively. Correction of the genomes using Illumina short reads was run by Pilon (Walker et al. 2014) and iCORN2 (Otto et al. 2010) through multiple iterations to eliminate errors from SMRT sequencing.

### Repeat annotation

RepeatModeler v1.0.11 (http://www.repeatmasker.org/RepeatModeler) was used to build a *de novo* repeats library, and then used RepeatMasker v 4.0.7 (http://www.repeatmasker.org) with search engine parameter as “ncbi”.

### Genome annotation

To develop open reading frame (ORF) in the new genome sequences, WebApollo 2.0 (Lee et al. 2013) was deployed with the genome sequence and the following tracks of evidence were added:

1. Gene prediction from COMPANION (Steinbiss et al. 2016) using *Trypanosoma brucei* as reference.
2. Gene prediction using AUGUSTUS (Stanke et al. 2004; Stanke and Morgenstern 2005) which was self-trained by CL Brener genome.
3. Annotation transfer from CL Brener by Exonerate (Slater and Birney 2005).
4. ESTs from available EST sequencing libraries in *T. cruzi* (retrieved from https://tritrypdb.org/tritrypdb/app/record/dataset/DS_6889a51dab).
5. Proteins from available Mass spec data for *T. cruzi* (Queiroz et al. 2013).
6. Strand-specific RNA-seq alignment data, the pipeline of which was followed as previously described (Kieft et al. 2020).

Each ORF along the genome was manually produced by the integration of all tracks.

InterProScan v5.31-70.0 (Jones et al. 2014) was used to detect protein families, domains and sites with all 11 default databases. Gene Ontology (GO) term was assigned also by InterProScan based on the protein domains results. Besides, BLASTP was used to search protein homology against *T. cruzi* CL Brener, *T. brucei, Leishmania major* databases from TriTrypDB release 39 (https://tritrypdb.org/tritrypdb/) and RefSeq non-redundant protein database, respectively, to determine the best hit for protein naming by in-house scripts. The parameter used for BLASTP was evalue <1e-10, identity >70% and coverage (length of alignment/length of target protein) >70%. Predicted pseudogenes were named by homology in RefSeq non-redundant nucleotide database with evalue <1e-30.

### Annotation of large gene families

A customized computational pipeline automated using PERL and Python scripts were developed for identifying members of large gene families in the genome. First, the annotated members of each gene family were searched against the Brazil A4 and Y C6 genome using BLASTN (version ncbi-blast-2.8.1+) with num_alignments and max_hsps arguments set to 100, the perc_identity argument set to 85. BLAST hits that have an overlap longer than 100 bp were merged if they match members from the same gene family. BLAST hits that were bracketed by longer hits from the same family were removed. A minimum length cutoff of 150 bp was applied to the BLAST hits. The remaining BLAST hits were considered new family member gene candidates. The new candidate genes were BLASTNed against all annotated transcripts from the genome of *T. cruzi* CL Brener strain (TriTrypDB release 34) (BLAST argument settings: num_alignments and max_hsps set to 50, perc_identity not set). Candidate genes were retained only if one of its top two best matches is a member of the candidate gene’s corresponding gene family.

Next, the boundaries of the candidate genes were refined by using model genes of each family in two steps. (1) Extending the candidate gene boundaries to include possible segments missed by previous steps. Using model gene sequences of each family to search the new genomes, and compare the coordinates of the matches to that of candidate genes. If > 50% overlap was found, and the non-overlapping length was < 1,000 bp, then the boundary of the candidate was extended according to the genomic match of the model gene. (2) The boundary of candidate genes was next subjected to small-scale trimmings. The candidate genes were BLASTed against model genes of the corresponding gene family (num_alignments and max_hsps arguments set to 100, the perc_identity argument set to 85). If a match was found within 100bp distance to the boundary of candidate genes, the candidate gene boundary was trimmed to match that of the model gene.

The start of mucin candidate genes was further refined using a conserved signal peptide sequence (in an alignment format allowing for minor variations). The signal peptide sequences were BLASTNed against mucin candidate genes (BLAST argument settings: num_alignments and max_hsps set to 200, perc_identity set to 65, gapopen and gapextend set to 1). Sequence upstream of signal peptide matches in the candidate genes was removed.

A final trim was applied to the boundaries of all candidate genes, as many of our BLAST steps could lead to inaccurate boundary identification due to 25% chance of a random matching an extra nucleotide base at the boundary and 6.25% chance for two extra bases and so forth, which could obscure start and stop codons. As an attempt to address this issue, we trimmed up to 10 bases which could reveal a start/stop codon that is in-frame with an existing stop/start codon.

Manual corrections of boundaries for gene family members were performed when necessary.

### Hi-C contact matrix

Hi-C contact matrix were analyzed by following the manual of <https://github.com/hms-dbmi/hic-data-analysis-bootcamp/, and then visualized in HiGlass (Kerpedjiev et al. 2018).

### Multidimensional scaling

K-tuple distance between genes are calculated with Clustal-Omega 1.2.4 (Sievers et al. 2011) using unaligned sequences with option parameters: “--full” and “--distmat-out”. Full alignment distance between genes are calculated with Clustal-Omega 1.2.4 using aligned sequences (aligned with Clustal-Omega 1.2.4 using default parameters) with options parameters: --full --full-iter --distmat-out. MDS is performed with the “cmdscale” function built-in R 3.6.3 with the input of a matrix of either pairwise K-tuple distances or full alignment distances. The results of MDS are visualized using the Shiny package 1.4.0.2 in R 3.6.3.

### Phylogenetic inference

Multiple sequence alignment was performed using MUSCLE (Edgar 2004). The resulting alignment was manually edited. Bayesian inference of phylogeny was performed using MrBayes v.3.2.6 (Ronquist et al. 2012) with the following parameters: nst = 6, rates = invgamma, Ngammacat = 8, Ngen = 10,000,000, nruns = 2, nchains = 4, and burn-infraction = 0.5. Convergence was determined by 25,000 post burn-in samples from two independent runs. The resulting phylogenetic tree was rendered in Figtree v.1.4.4. Node support values are given in percent posterior probability.

## Data access

All raw and processed sequencing data generated in this study have been submitted to the NCBI Sequence Read Archive (SRA; https://www.ncbi.nlm.nih.gov/sra/) under accession number SRR118039885-SRR118039888 (Brazil A4) and SRR11845028-SRR11845031 (Y C6). The genome and annotation data generated in this study have been submitted to the NCBI BioProject database (https://www.ncbi.nlm.nih.gov/bioproject/) under accession number PRJNA512864 (Brazil A4) and PRJNA554625 (Y C6). The assembled and annotated genomes are also accessible in the TritrypDB database (https://tritrypdb.org/tritrypdb/).

## Acknowledgements

We thank Dr. Todd Minning for initial contributions this project, Dr. Robert Sabatini from the University of Georgia for insightful discussions, and Dr. Benedikt Brink from Ludwig-Maximilians-Universität for advice and for testing our data in his genome phasing pipeline. This work was supported by funding from the National Institutes of Health, (USA) grants R03 AI124228 and R01 AI124692 to RLT.

