## Supplemental Figure and Tables for "Strain-specific genome evolution in *Trypanosoma cruzi*, the agent of Chagas disease"

### Slide 1
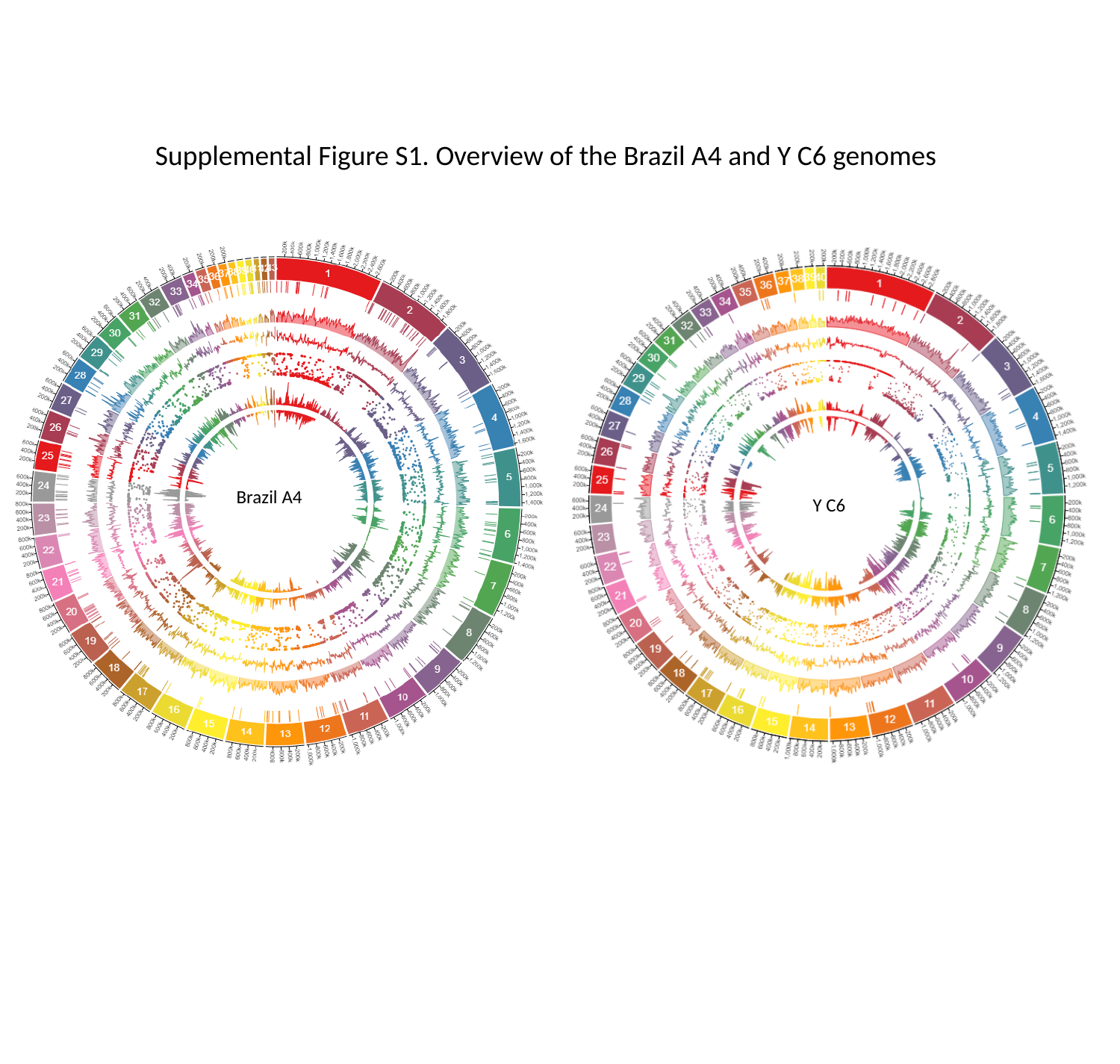

#### Supplemental Figure S1. Overview of the Brazil A4 and Y C6 genomes
Brazil A4
Y C6

### Slide 2
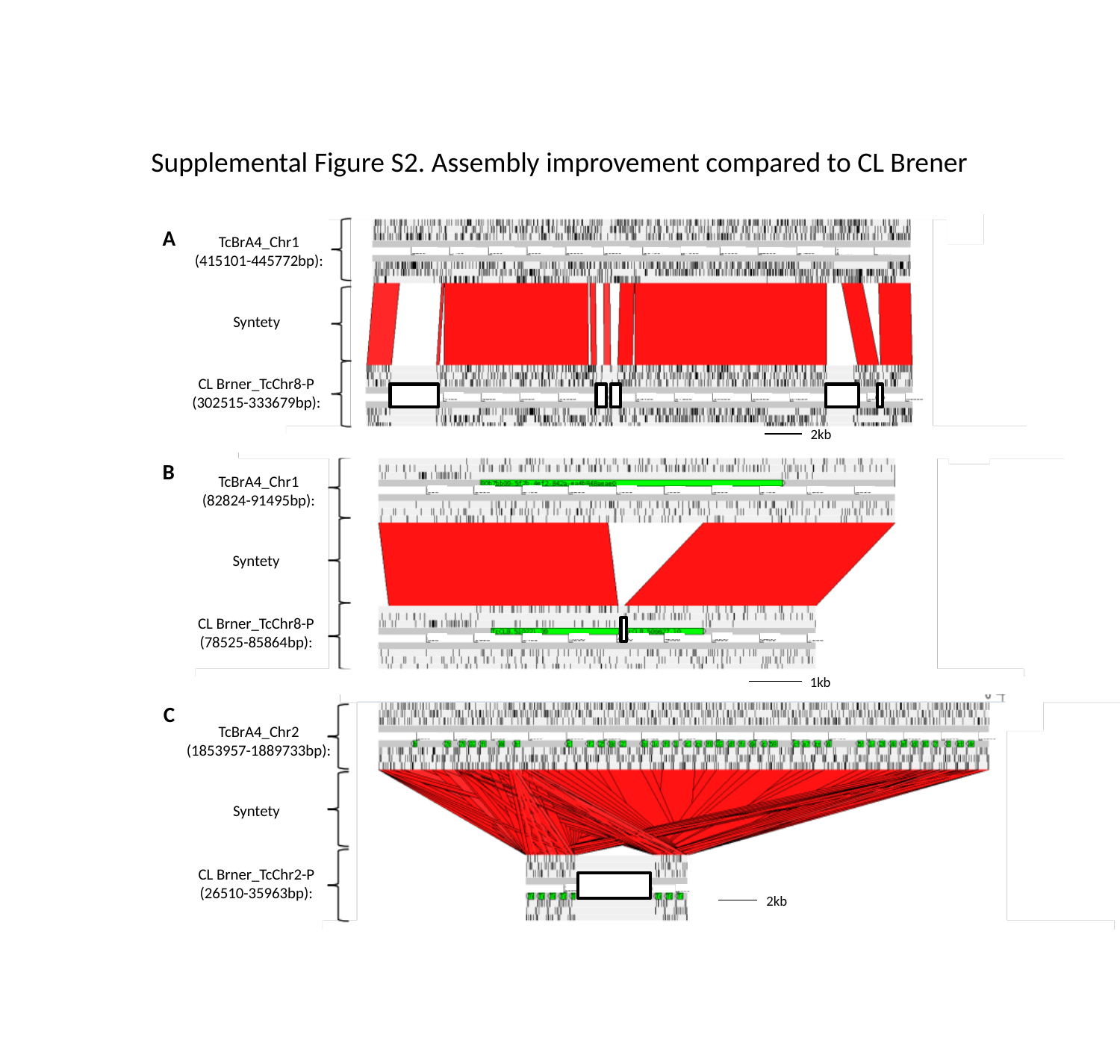

Supplemental Figure S2. Assembly improvement compared to CL Brener
A
TcBrA4_Chr1
(415101-445772bp):
Syntety
CL Brner_TcChr8-P
(302515-333679bp):
2kb
TcBrA4_Chr1
(82824-91495bp):
Syntety
CL Brner_TcChr8-P
(78525-85864bp):
1kb
B
C
TcBrA4_Chr2
(1853957-1889733bp):
Syntety
2kb
CL Brner_TcChr2-P
(26510-35963bp):

### Slide 3
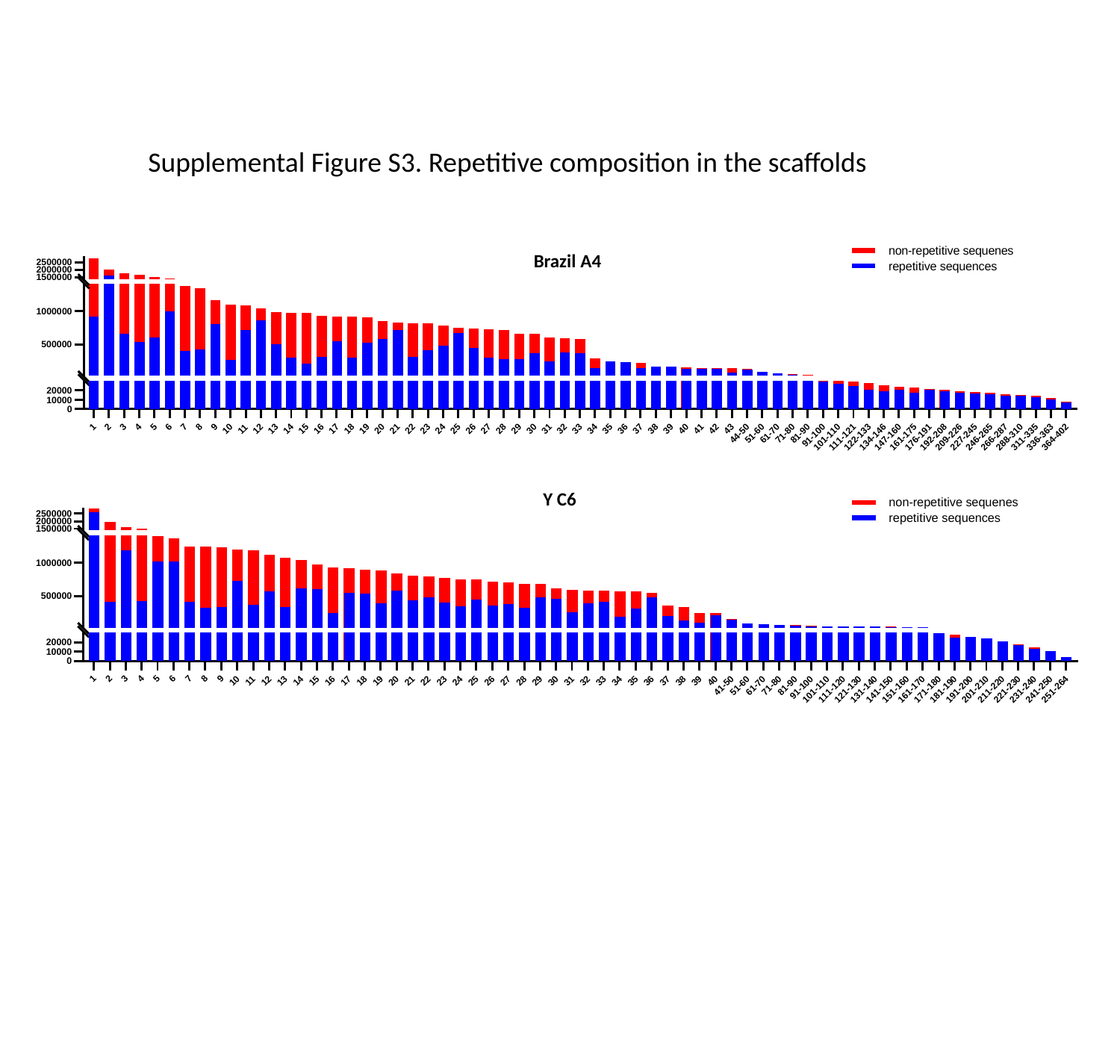

Supplemental Figure S3. Repetitive composition in the scaffolds
Brazil A4
Y C6

### Slide 4
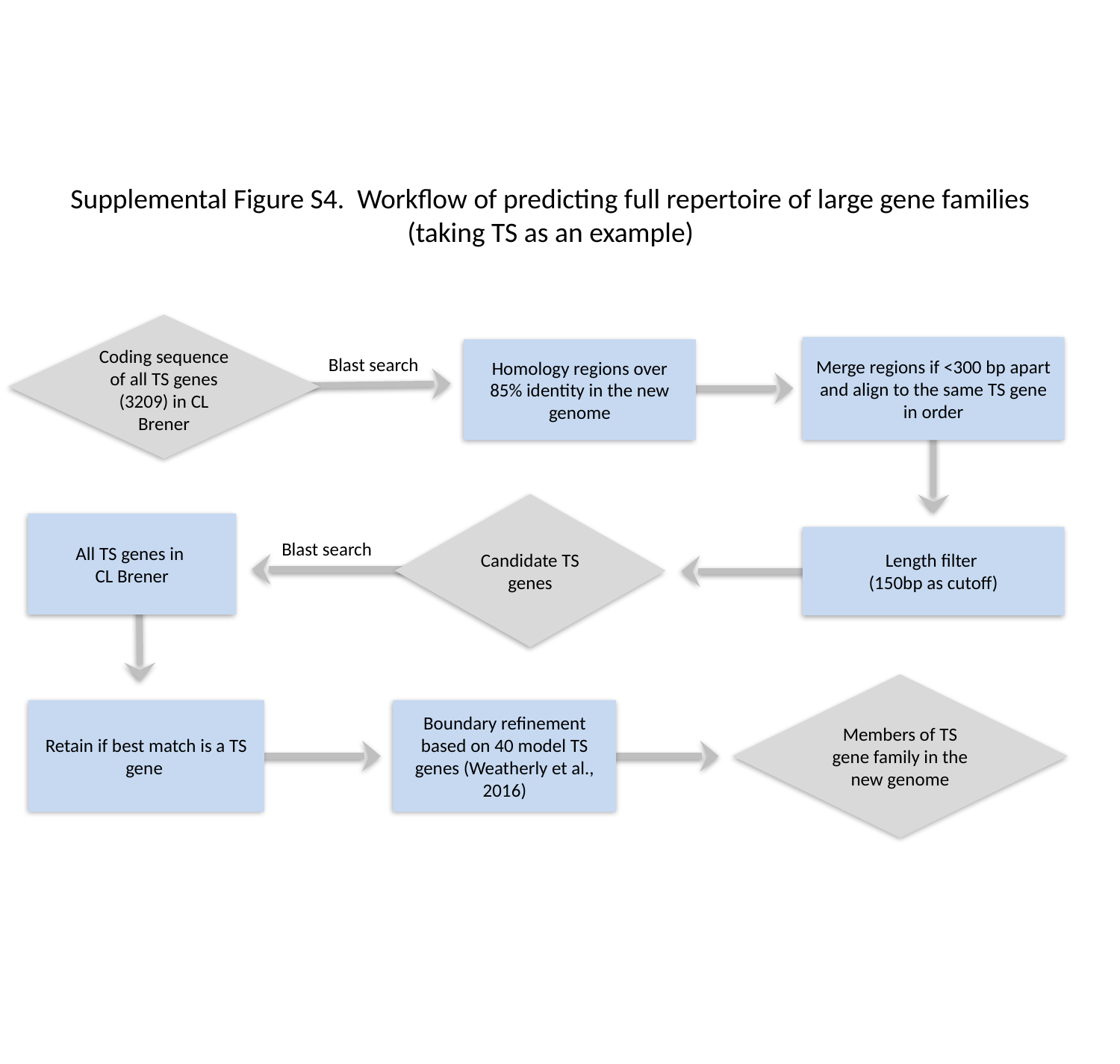

Supplemental Figure S4. Workflow of predicting full repertoire of large gene families (taking TS as an example)
Coding sequence of all TS genes (3209) in CL Brener
Merge regions if <300 bp apart and align to the same TS gene in order
Homology regions over 85% identity in the new genome
Blast search
Candidate TS genes
All TS genes in
CL Brener
Length filter
(150bp as cutoff)
Blast search
Members of TS gene family in the new genome
Boundary refinement based on 40 model TS genes (Weatherly et al., 2016)
Retain if best match is a TS gene

### Slide 5
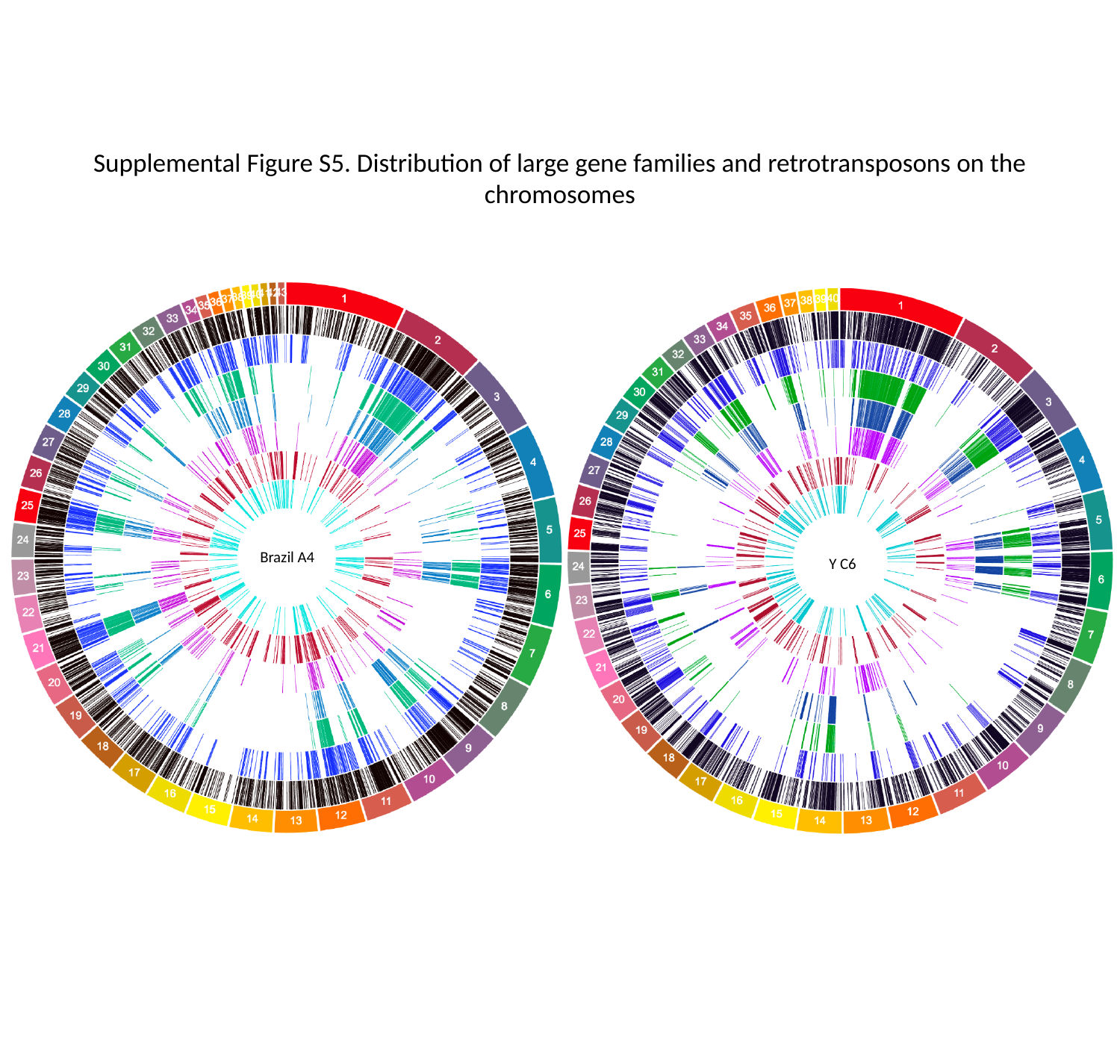

Supplemental Figure S5. Distribution of large gene families and retrotransposons on the chromosomes
Brazil A4
Y C6

### Slide 6
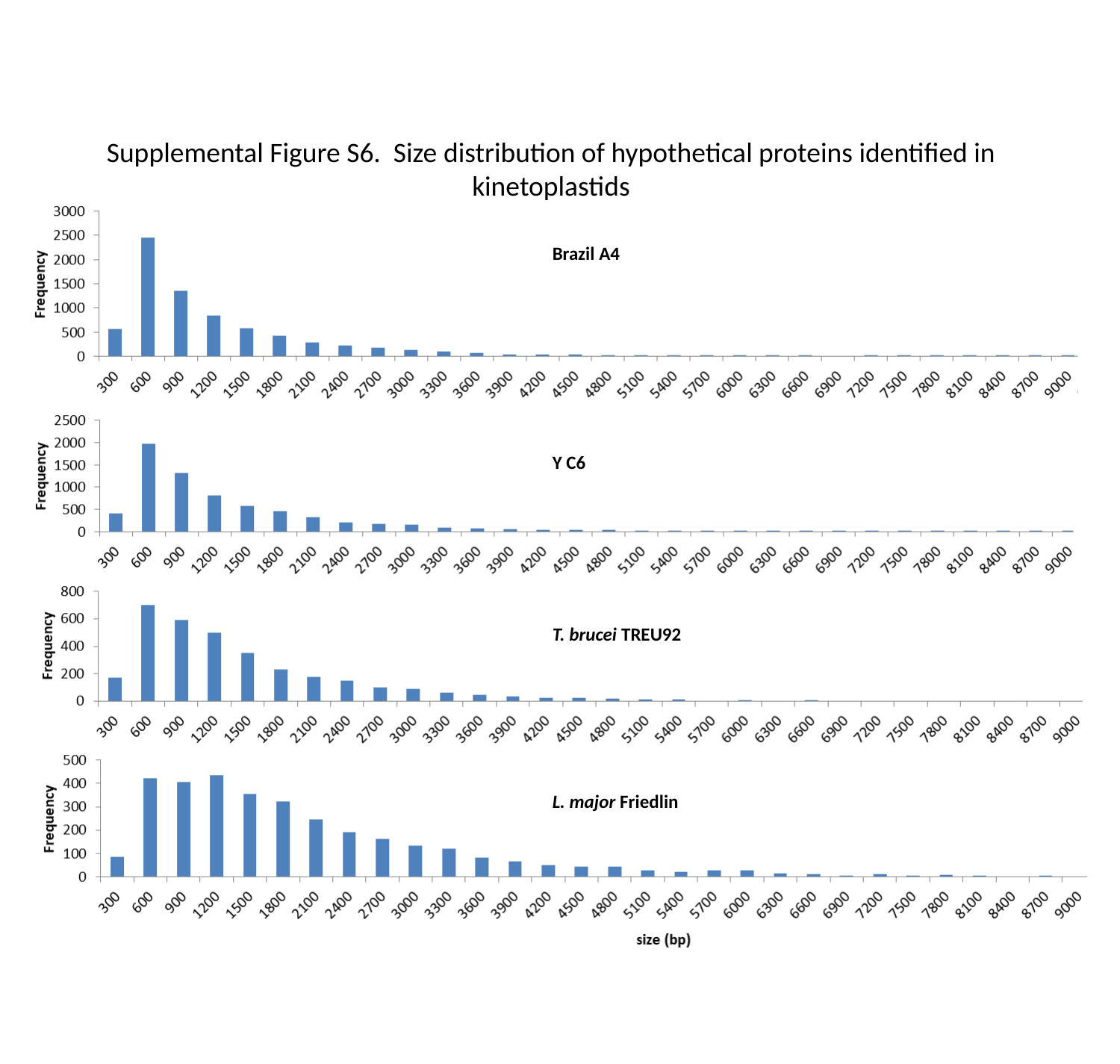

Supplemental Figure S6. Size distribution of hypothetical proteins identified in kinetoplastids
Brazil A4
Y C6
T. brucei TREU92
L. major Friedlin

### Slide 7
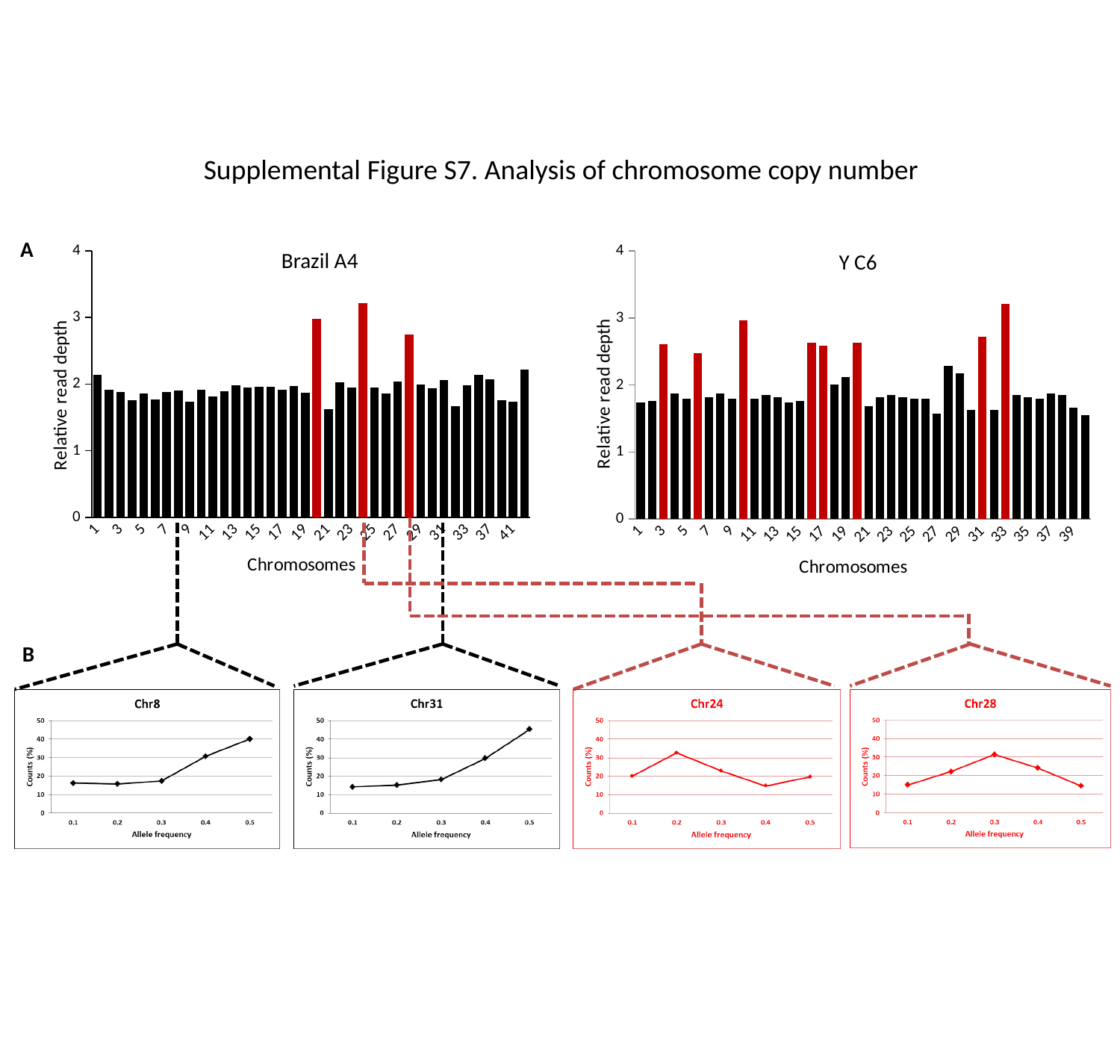

Supplemental Figure S7. Analysis of chromosome copy number
###### Chart
| Category |
|---|
###### Chart
| Category | |
|---|---|
| 1.0 | 2.137546941677378 |
| 2.0 | 1.916903904292178 |
| 3.0 | 1.875891526662686 |
| 4.0 | 1.757709688543571 |
| 5.0 | 1.864411314833684 |
| 6.0 | 1.767899461151672 |
| 7.0 | 1.884991372719802 |
| 8.0 | 1.903554958015068 |
| 9.0 | 1.738303635401376 |
| 10.0 | 1.916255454608581 |
| 11.0 | 1.81865396983294 |
| 12.0 | 1.88933800678747 |
| 13.0 | 1.978887397988347 |
| 14.0 | 1.95161565338353 |
| 15.0 | 1.9607223406364 |
| 16.0 | 1.95671325384198 |
| 17.0 | 1.91625707036528 |
| 18.0 | 1.973503700163723 |
| 19.0 | 1.871245059959524 |
| 20.0 | 2.979942171113132 |
| 21.0 | 1.62182070033172 |
| 22.0 | 2.021815059002107 |
| 23.0 | 1.943872681244968 |
| 24.0 | 3.216127897253692 |
| 25.0 | 1.944145106918646 |
| 26.0 | 1.853915335241648 |
| 27.0 | 2.03766577028157 |
| 28.0 | 2.744463974811246 |
| 29.0 | 1.990947454138174 |
| 30.0 | 1.934207211089682 |
| 31.0 | 2.060785351015346 |
| 32.0 | 1.665652816186945 |
| 33.0 | 1.986906525435337 |
| 34.0 | 2.136259840137004 |
| 37.0 | 2.068529730691357 |
| 40.0 | 1.758761003591471 |
| 41.0 | 1.740870857376202 |
| 43.0 | 2.212905803274534 |A
Brazil A4
Y C6
B

### Slide 8
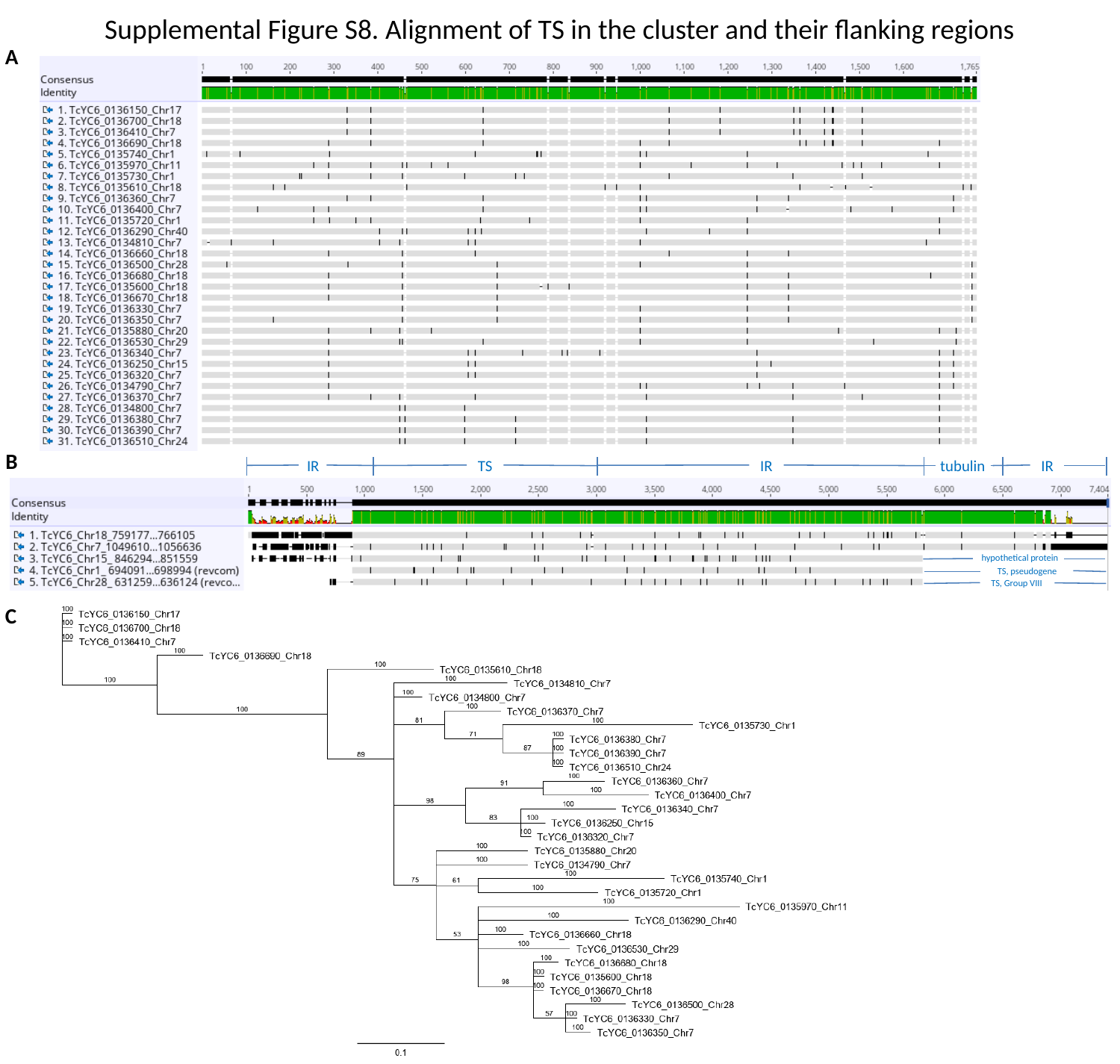

Supplemental Figure S8. Alignment of TS in the cluster and their flanking regions
A
B
IR
TS
IR
tubulin
IR
hypothetical protein
 TS, pseudogene
TS, Group VIII
C

### Slide 9
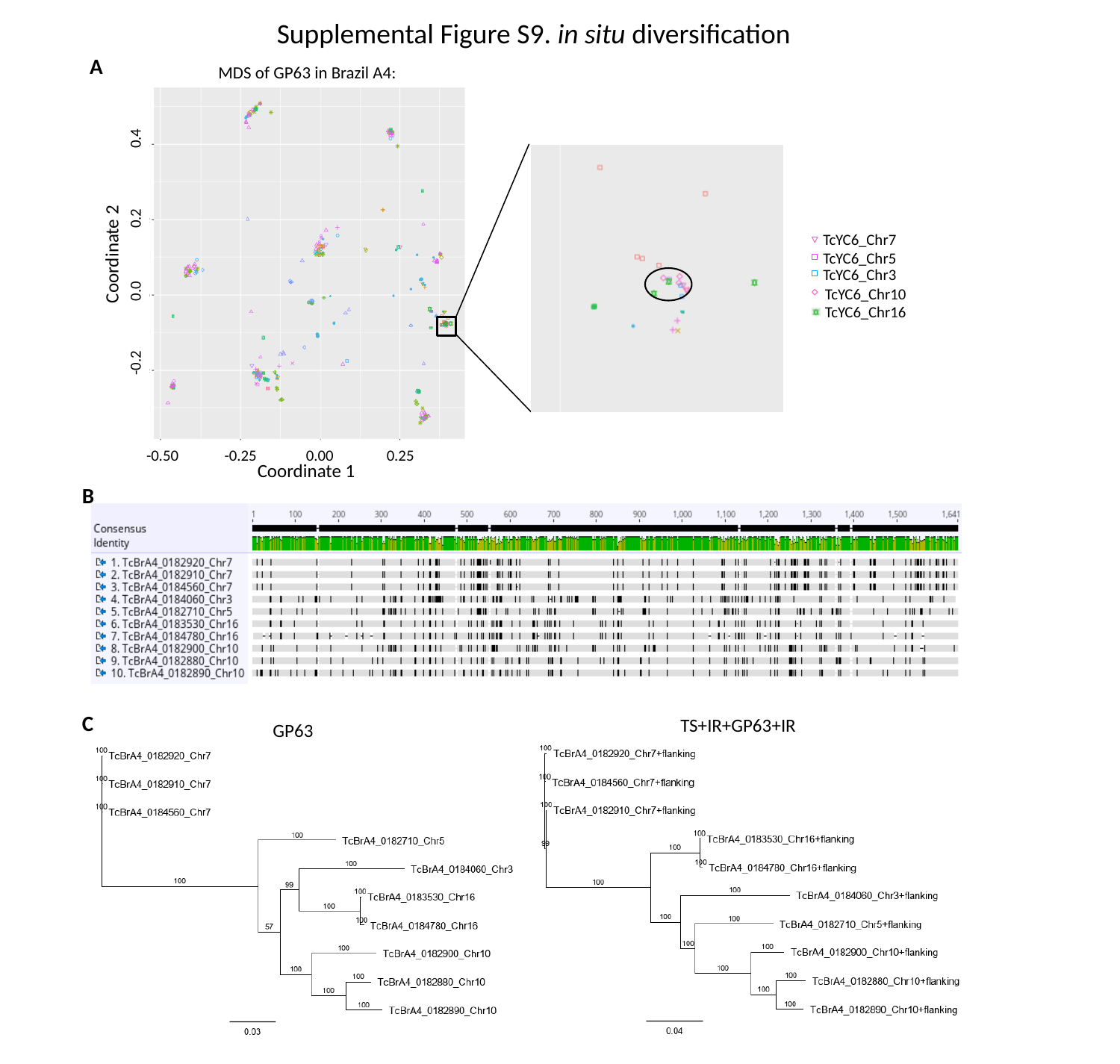

#### Supplemental Figure S9. in situ diversification
A
MDS of GP63 in Brazil A4:
TcYC6_Chr7
Coordinate 2
 -0.2 0.0 0.2 0.4
TcYC6_Chr5
TcYC6_Chr3
TcYC6_Chr10
TcYC6_Chr16
 -0.50 -0.25 0.00 0.25
Coordinate 1
B
C
TS+IR+GP63+IR
GP63

### Slide 10
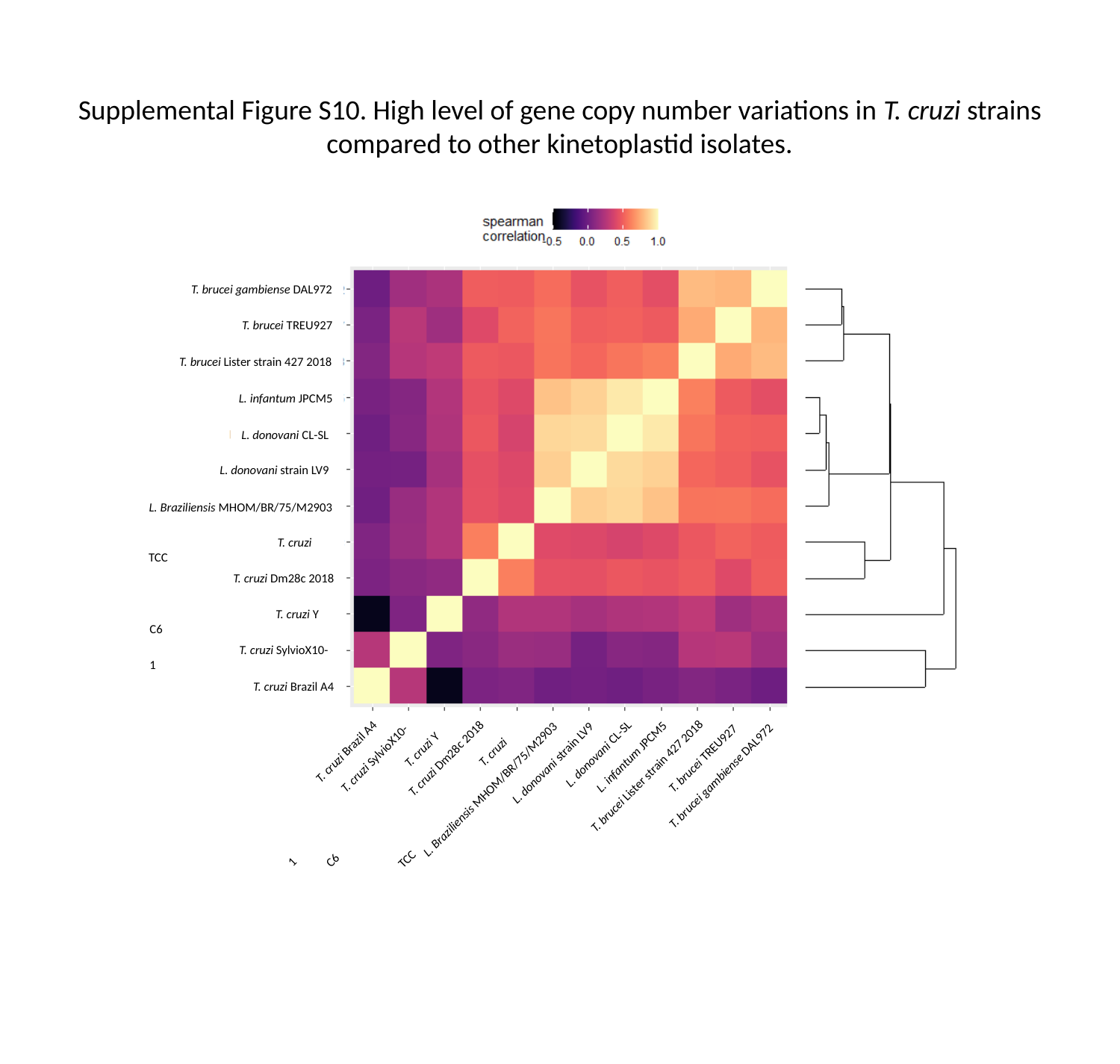

#### Supplemental Figure S10. High level of gene copy number variations in T. cruzi strains compared to other kinetoplastid isolates.
T. brucei gambiense DAL972
T. brucei TREU927
T. brucei Lister strain 427 2018
L. infantum JPCM5
L. donovani CL-SL
L. donovani strain LV9
L. Braziliensis MHOM/BR/75/M2903
 T. cruzi TCC
 T. cruzi Dm28c 2018
 T. cruzi Y C6
 T. cruzi SylvioX10-1
 T. cruzi Brazil A4
L. donovani CL-SL
L. infantum JPCM5
T. brucei TREU927
L. donovani strain LV9
T. brucei Lister strain 427 2018
T. brucei gambiense DAL972
 T. cruzi Brazil A4
 T. cruzi SylvioX10-1
 T. cruzi Dm28c 2018
 T. cruzi Y C6
L. Braziliensis MHOM/BR/75/M2903
 T. cruzi TCC

### Slide 11
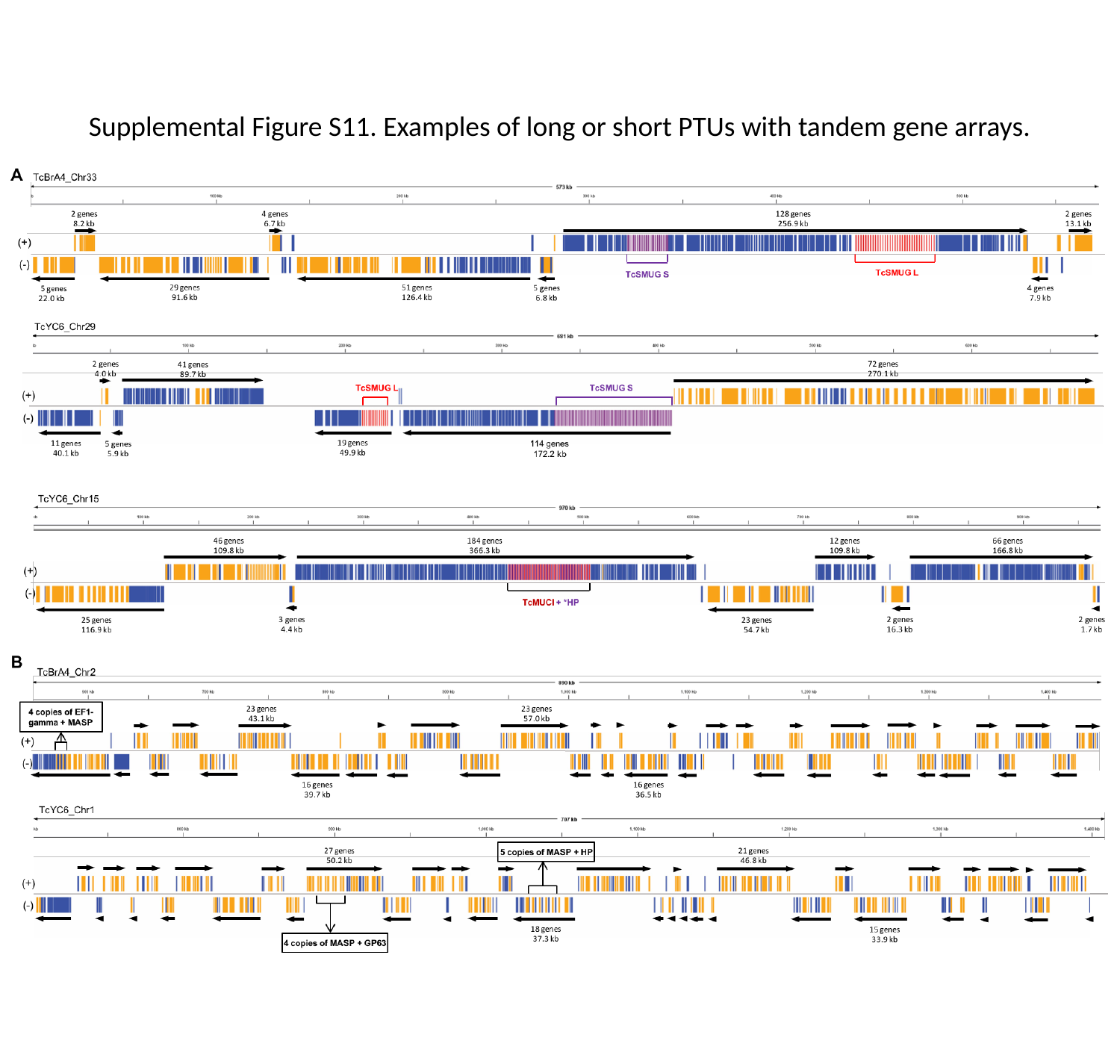

#### Supplemental Figure S11. Examples of long or short PTUs with tandem gene arrays.

### Slide 12
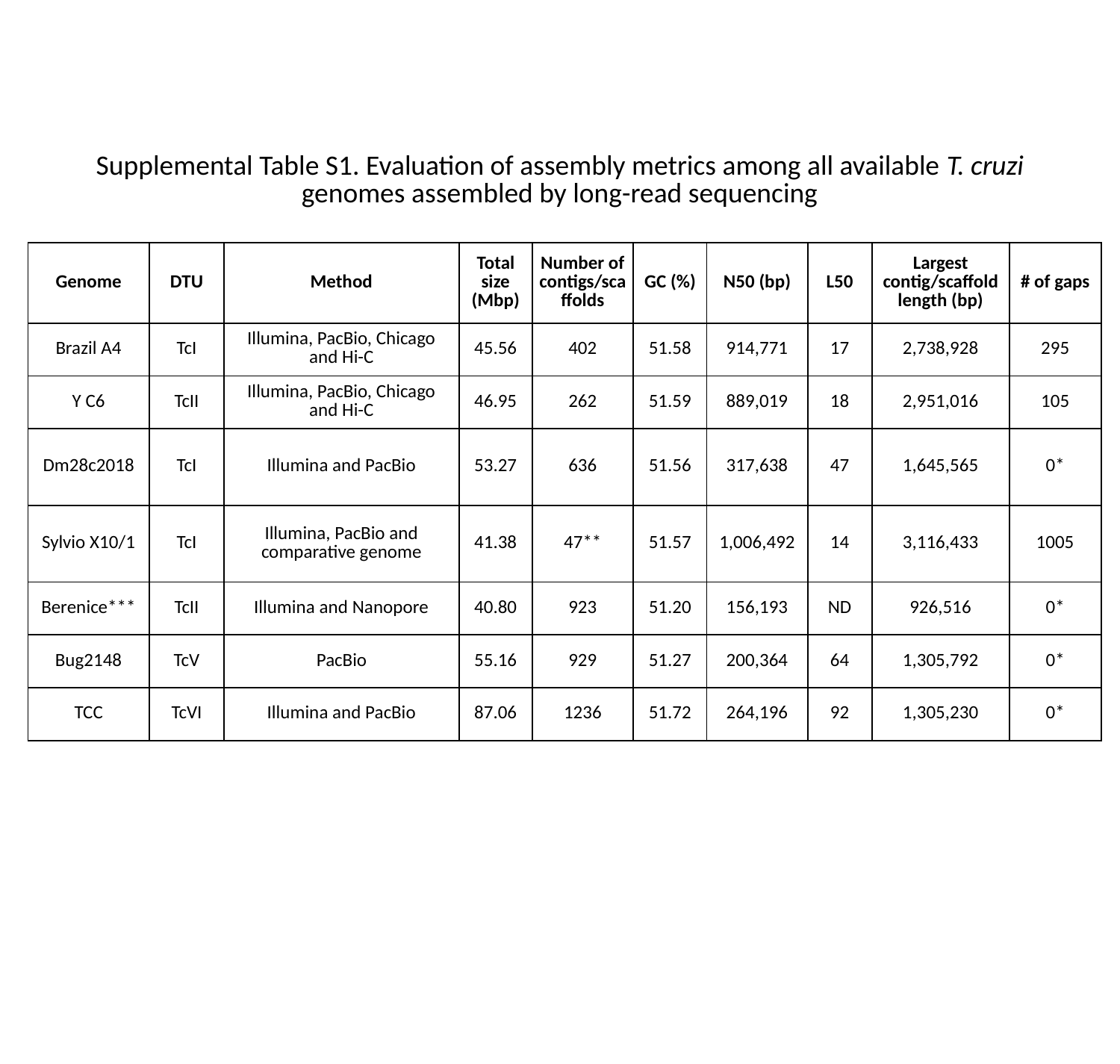

#### Supplemental Table S1. Evaluation of assembly metrics among all available T. cruzi genomes assembled by long-read sequencing
| Genome | DTU | Method | Total size (Mbp) | Number of contigs/scaffolds | GC (%) | N50 (bp) | L50 | Largest contig/scaffold length (bp) | # of gaps |
| --- | --- | --- | --- | --- | --- | --- | --- | --- | --- |
| Brazil A4 | TcI | Illumina, PacBio, Chicago and Hi-C | 45.56 | 402 | 51.58 | 914,771 | 17 | 2,738,928 | 295 |
| Y C6 | TcII | Illumina, PacBio, Chicago and Hi-C | 46.95 | 262 | 51.59 | 889,019 | 18 | 2,951,016 | 105 |
| Dm28c2018 | TcI | Illumina and PacBio | 53.27 | 636 | 51.56 | 317,638 | 47 | 1,645,565 | 0\* |
| Sylvio X10/1 | TcI | Illumina, PacBio and comparative genome | 41.38 | 47\*\* | 51.57 | 1,006,492 | 14 | 3,116,433 | 1005 |
| Berenice\*\*\* | TcII | Illumina and Nanopore | 40.80 | 923 | 51.20 | 156,193 | ND | 926,516 | 0\* |
| Bug2148 | TcV | PacBio | 55.16 | 929 | 51.27 | 200,364 | 64 | 1,305,792 | 0\* |
| TCC | TcVI | Illumina and PacBio | 87.06 | 1236 | 51.72 | 264,196 | 92 | 1,305,230 | 0\* |

### Slide 13
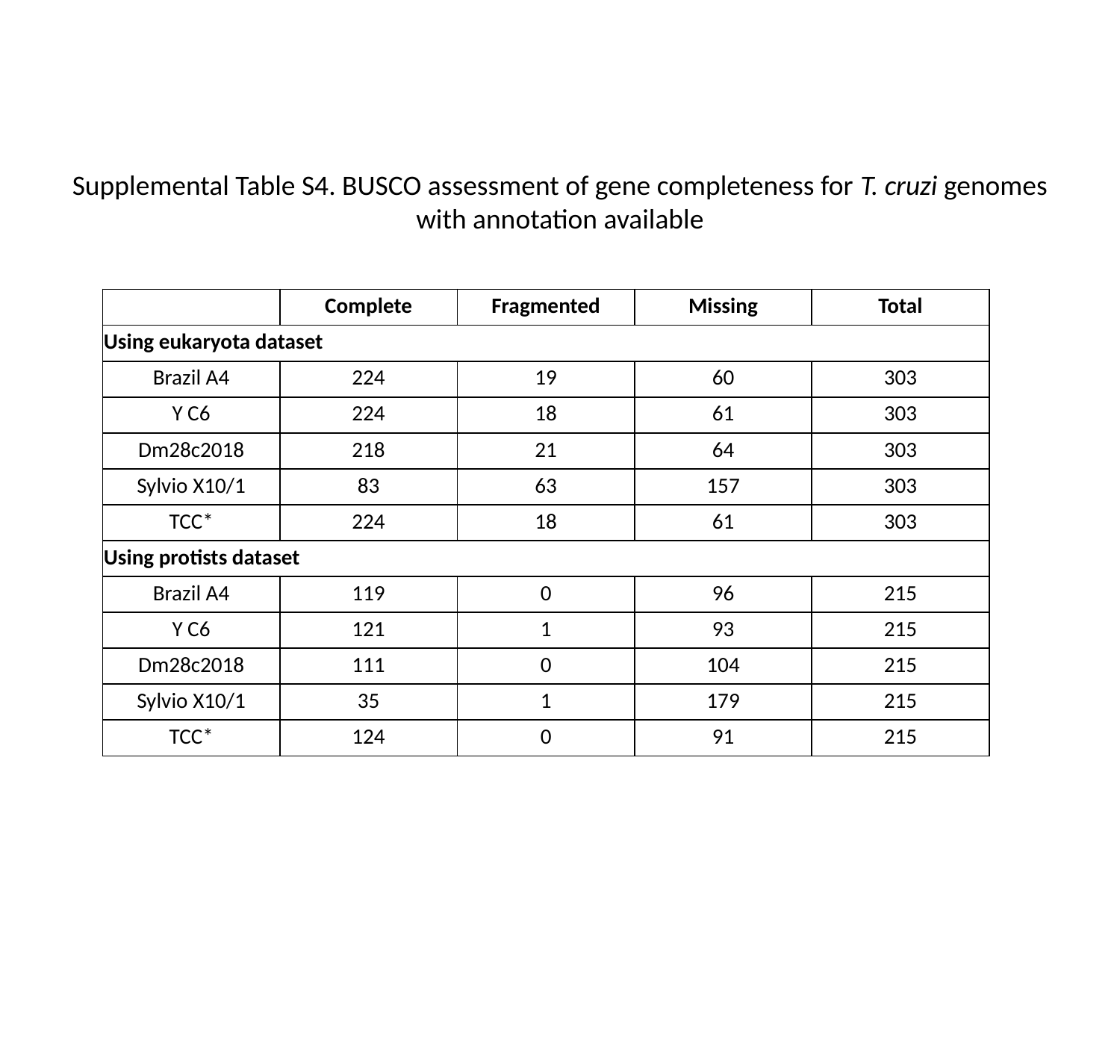

#### Supplemental Table S4. BUSCO assessment of gene completeness for T. cruzi genomes with annotation available
| | Complete | Fragmented | Missing | Total |
| --- | --- | --- | --- | --- |
| Using eukaryota dataset | | | | |
| Brazil A4 | 224 | 19 | 60 | 303 |
| Y C6 | 224 | 18 | 61 | 303 |
| Dm28c2018 | 218 | 21 | 64 | 303 |
| Sylvio X10/1 | 83 | 63 | 157 | 303 |
| TCC\* | 224 | 18 | 61 | 303 |
| Using protists dataset | | | | |
| Brazil A4 | 119 | 0 | 96 | 215 |
| Y C6 | 121 | 1 | 93 | 215 |
| Dm28c2018 | 111 | 0 | 104 | 215 |
| Sylvio X10/1 | 35 | 1 | 179 | 215 |
| TCC\* | 124 | 0 | 91 | 215 |

### Slide 14
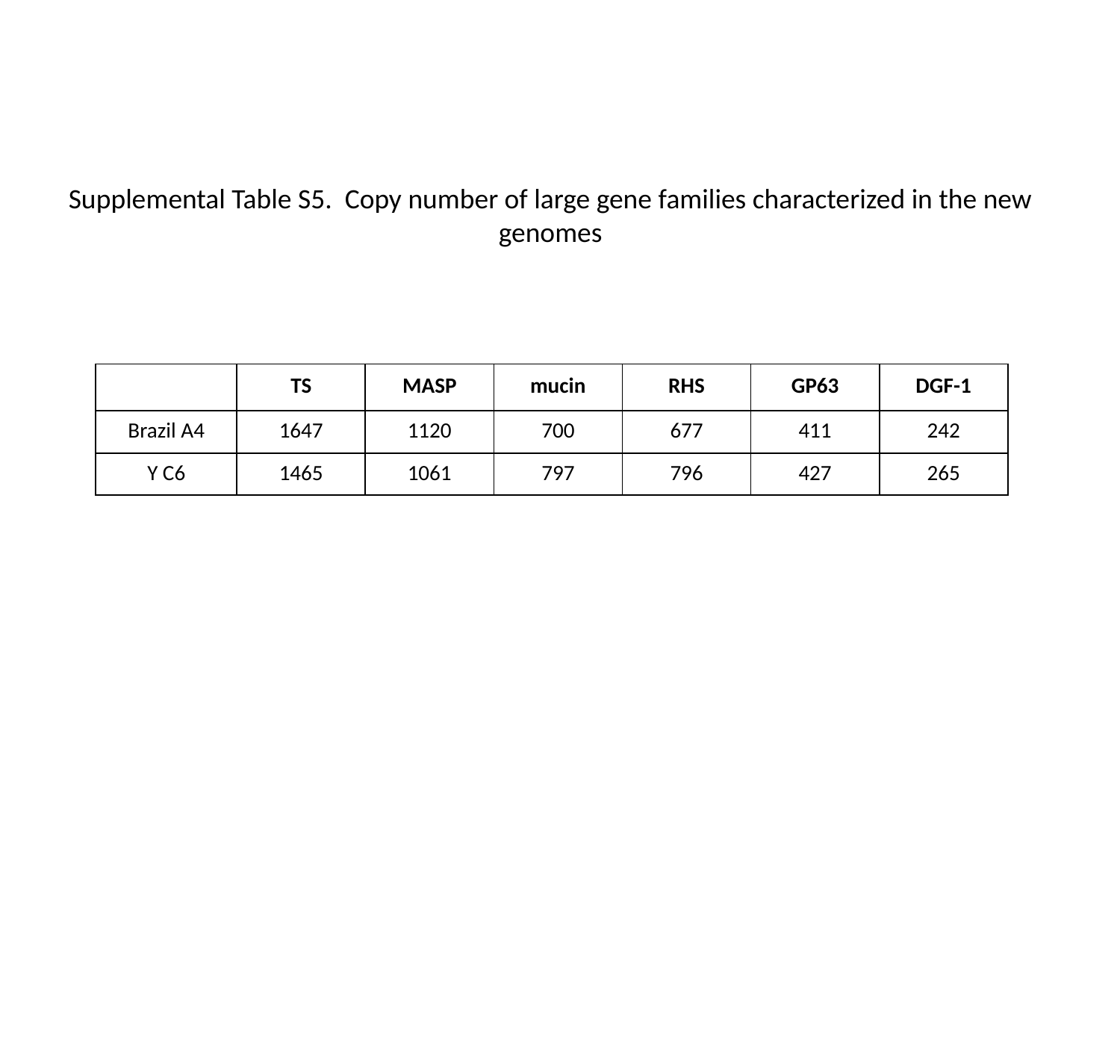

Supplemental Table S5. Copy number of large gene families characterized in the new genomes
| | TS | MASP | mucin | RHS | GP63 | DGF-1 |
| --- | --- | --- | --- | --- | --- | --- |
| Brazil A4 | 1647 | 1120 | 700 | 677 | 411 | 242 |
| Y C6 | 1465 | 1061 | 797 | 796 | 427 | 265 |

### Slide 15
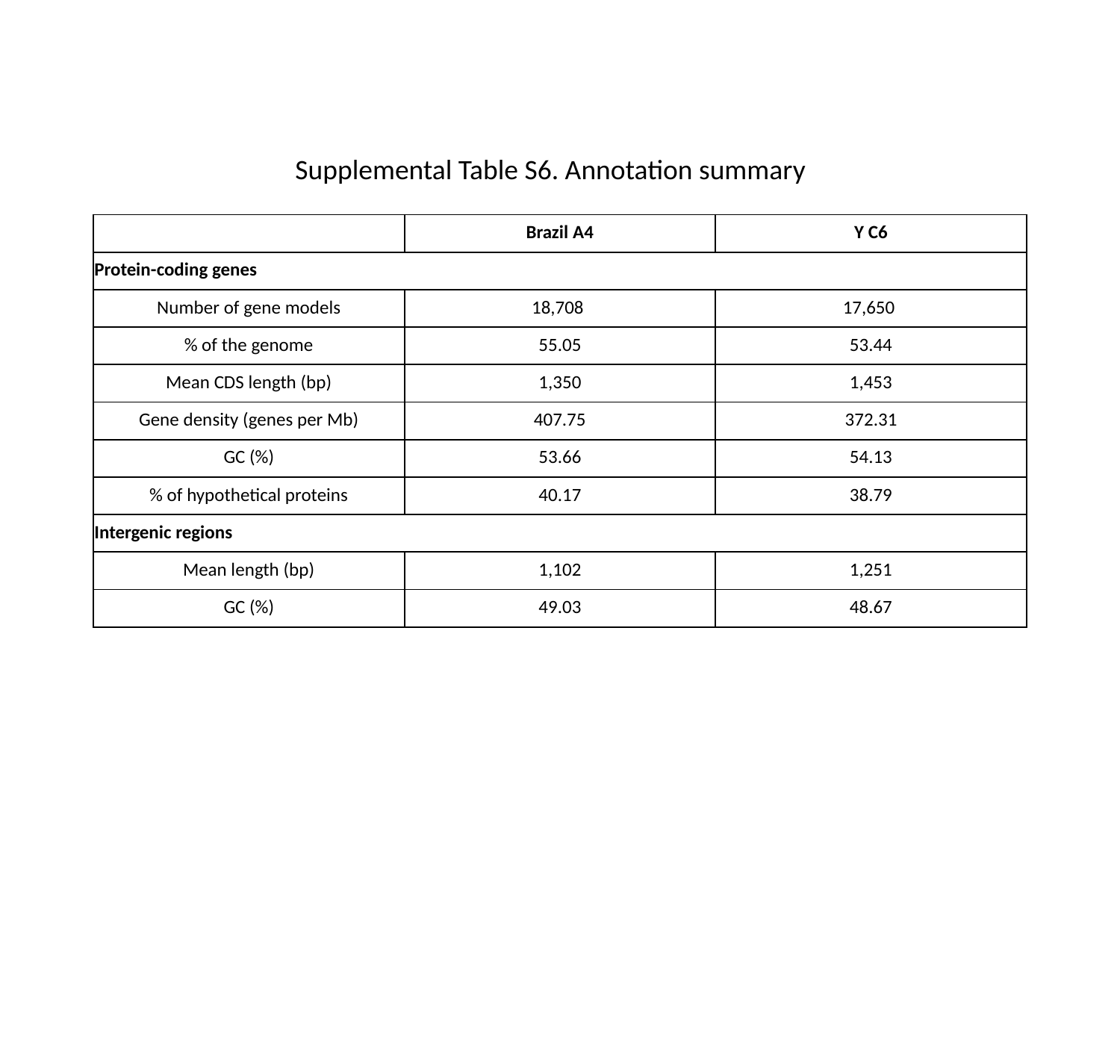

Supplemental Table S6. Annotation summary
| | Brazil A4 | Y C6 |
| --- | --- | --- |
| Protein-coding genes | | |
| Number of gene models | 18,708 | 17,650 |
| % of the genome | 55.05 | 53.44 |
| Mean CDS length (bp) | 1,350 | 1,453 |
| Gene density (genes per Mb) | 407.75 | 372.31 |
| GC (%) | 53.66 | 54.13 |
| % of hypothetical proteins | 40.17 | 38.79 |
| Intergenic regions | | |
| Mean length (bp) | 1,102 | 1,251 |
| GC (%) | 49.03 | 48.67 |

### Slide 16
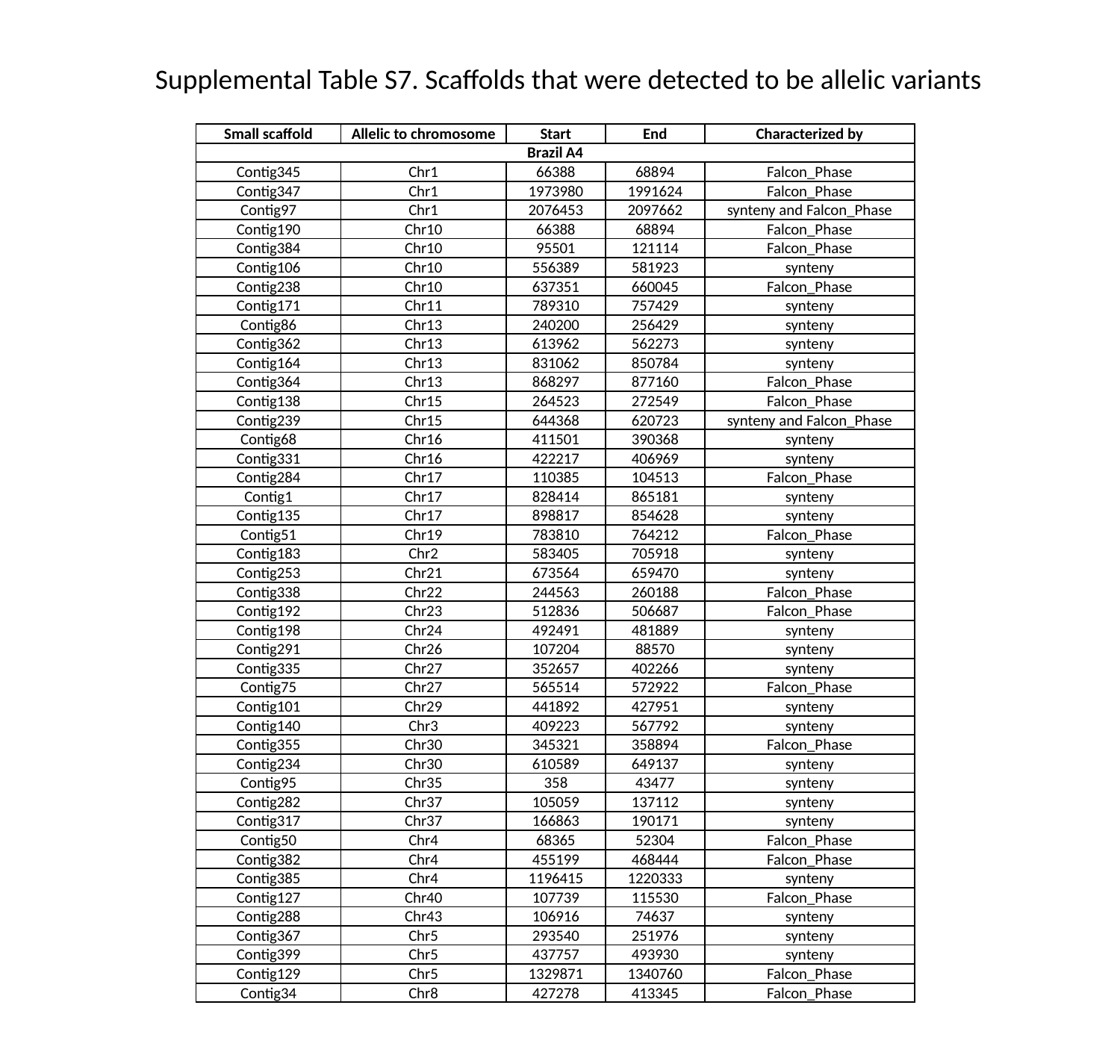

Supplemental Table S7. Scaffolds that were detected to be allelic variants
| Small scaffold | Allelic to chromosome | Start | End | Characterized by |
| --- | --- | --- | --- | --- |
| Brazil A4 | | | | |
| Contig345 | Chr1 | 66388 | 68894 | Falcon\_Phase |
| Contig347 | Chr1 | 1973980 | 1991624 | Falcon\_Phase |
| Contig97 | Chr1 | 2076453 | 2097662 | synteny and Falcon\_Phase |
| Contig190 | Chr10 | 66388 | 68894 | Falcon\_Phase |
| Contig384 | Chr10 | 95501 | 121114 | Falcon\_Phase |
| Contig106 | Chr10 | 556389 | 581923 | synteny |
| Contig238 | Chr10 | 637351 | 660045 | Falcon\_Phase |
| Contig171 | Chr11 | 789310 | 757429 | synteny |
| Contig86 | Chr13 | 240200 | 256429 | synteny |
| Contig362 | Chr13 | 613962 | 562273 | synteny |
| Contig164 | Chr13 | 831062 | 850784 | synteny |
| Contig364 | Chr13 | 868297 | 877160 | Falcon\_Phase |
| Contig138 | Chr15 | 264523 | 272549 | Falcon\_Phase |
| Contig239 | Chr15 | 644368 | 620723 | synteny and Falcon\_Phase |
| Contig68 | Chr16 | 411501 | 390368 | synteny |
| Contig331 | Chr16 | 422217 | 406969 | synteny |
| Contig284 | Chr17 | 110385 | 104513 | Falcon\_Phase |
| Contig1 | Chr17 | 828414 | 865181 | synteny |
| Contig135 | Chr17 | 898817 | 854628 | synteny |
| Contig51 | Chr19 | 783810 | 764212 | Falcon\_Phase |
| Contig183 | Chr2 | 583405 | 705918 | synteny |
| Contig253 | Chr21 | 673564 | 659470 | synteny |
| Contig338 | Chr22 | 244563 | 260188 | Falcon\_Phase |
| Contig192 | Chr23 | 512836 | 506687 | Falcon\_Phase |
| Contig198 | Chr24 | 492491 | 481889 | synteny |
| Contig291 | Chr26 | 107204 | 88570 | synteny |
| Contig335 | Chr27 | 352657 | 402266 | synteny |
| Contig75 | Chr27 | 565514 | 572922 | Falcon\_Phase |
| Contig101 | Chr29 | 441892 | 427951 | synteny |
| Contig140 | Chr3 | 409223 | 567792 | synteny |
| Contig355 | Chr30 | 345321 | 358894 | Falcon\_Phase |
| Contig234 | Chr30 | 610589 | 649137 | synteny |
| Contig95 | Chr35 | 358 | 43477 | synteny |
| Contig282 | Chr37 | 105059 | 137112 | synteny |
| Contig317 | Chr37 | 166863 | 190171 | synteny |
| Contig50 | Chr4 | 68365 | 52304 | Falcon\_Phase |
| Contig382 | Chr4 | 455199 | 468444 | Falcon\_Phase |
| Contig385 | Chr4 | 1196415 | 1220333 | synteny |
| Contig127 | Chr40 | 107739 | 115530 | Falcon\_Phase |
| Contig288 | Chr43 | 106916 | 74637 | synteny |
| Contig367 | Chr5 | 293540 | 251976 | synteny |
| Contig399 | Chr5 | 437757 | 493930 | synteny |
| Contig129 | Chr5 | 1329871 | 1340760 | Falcon\_Phase |
| Contig34 | Chr8 | 427278 | 413345 | Falcon\_Phase |

### Slide 17
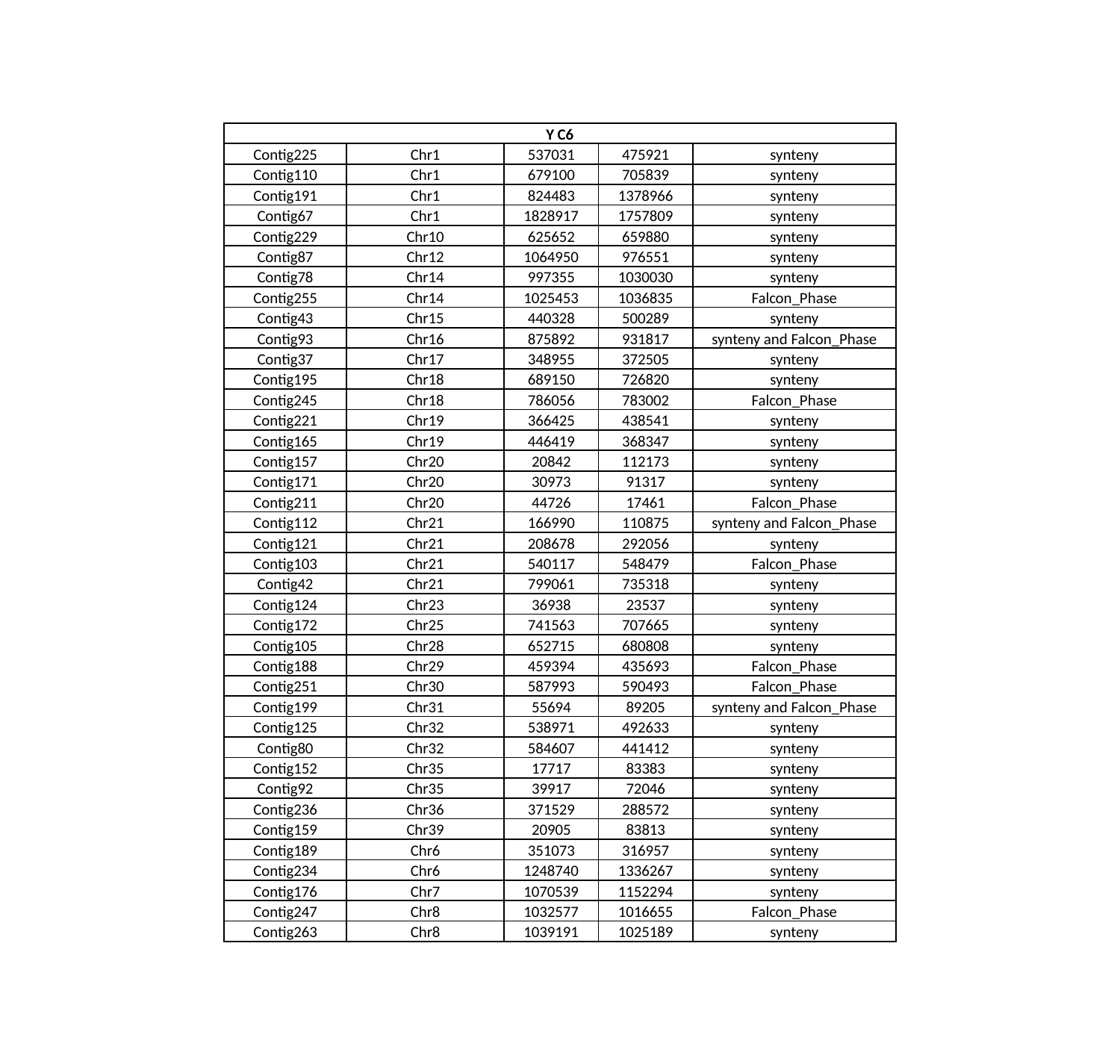

| Y C6 | | | | |
| --- | --- | --- | --- | --- |
| Contig225 | Chr1 | 537031 | 475921 | synteny |
| Contig110 | Chr1 | 679100 | 705839 | synteny |
| Contig191 | Chr1 | 824483 | 1378966 | synteny |
| Contig67 | Chr1 | 1828917 | 1757809 | synteny |
| Contig229 | Chr10 | 625652 | 659880 | synteny |
| Contig87 | Chr12 | 1064950 | 976551 | synteny |
| Contig78 | Chr14 | 997355 | 1030030 | synteny |
| Contig255 | Chr14 | 1025453 | 1036835 | Falcon\_Phase |
| Contig43 | Chr15 | 440328 | 500289 | synteny |
| Contig93 | Chr16 | 875892 | 931817 | synteny and Falcon\_Phase |
| Contig37 | Chr17 | 348955 | 372505 | synteny |
| Contig195 | Chr18 | 689150 | 726820 | synteny |
| Contig245 | Chr18 | 786056 | 783002 | Falcon\_Phase |
| Contig221 | Chr19 | 366425 | 438541 | synteny |
| Contig165 | Chr19 | 446419 | 368347 | synteny |
| Contig157 | Chr20 | 20842 | 112173 | synteny |
| Contig171 | Chr20 | 30973 | 91317 | synteny |
| Contig211 | Chr20 | 44726 | 17461 | Falcon\_Phase |
| Contig112 | Chr21 | 166990 | 110875 | synteny and Falcon\_Phase |
| Contig121 | Chr21 | 208678 | 292056 | synteny |
| Contig103 | Chr21 | 540117 | 548479 | Falcon\_Phase |
| Contig42 | Chr21 | 799061 | 735318 | synteny |
| Contig124 | Chr23 | 36938 | 23537 | synteny |
| Contig172 | Chr25 | 741563 | 707665 | synteny |
| Contig105 | Chr28 | 652715 | 680808 | synteny |
| Contig188 | Chr29 | 459394 | 435693 | Falcon\_Phase |
| Contig251 | Chr30 | 587993 | 590493 | Falcon\_Phase |
| Contig199 | Chr31 | 55694 | 89205 | synteny and Falcon\_Phase |
| Contig125 | Chr32 | 538971 | 492633 | synteny |
| Contig80 | Chr32 | 584607 | 441412 | synteny |
| Contig152 | Chr35 | 17717 | 83383 | synteny |
| Contig92 | Chr35 | 39917 | 72046 | synteny |
| Contig236 | Chr36 | 371529 | 288572 | synteny |
| Contig159 | Chr39 | 20905 | 83813 | synteny |
| Contig189 | Chr6 | 351073 | 316957 | synteny |
| Contig234 | Chr6 | 1248740 | 1336267 | synteny |
| Contig176 | Chr7 | 1070539 | 1152294 | synteny |
| Contig247 | Chr8 | 1032577 | 1016655 | Falcon\_Phase |
| Contig263 | Chr8 | 1039191 | 1025189 | synteny |
